# Mechanisms of USP18 deISGylation revealed by comparative analysis with its human paralog USP41

**DOI:** 10.1101/2024.05.28.596309

**Authors:** Thomas Bonacci, Derek L Bolhuis, Nicholas G Brown, Michael J Emanuele

## Abstract

The ubiquitin-like protein ISG15 (interferon-<u>s</u>timulated <u>g</u>ene <u>15</u>) regulates the host response to bacterial and viral infections through its conjugation to proteins (ISGylation) following interferon production. ISGylation is antagonized by the highly specific cysteine protease USP18, which is the major deISGylating enzyme. However, mechanisms underlying USP18’s extraordinary specificity towards ISG15 remains elusive. Here, we show that USP18 interacts with its paralog USP41, whose catalytic domain shares 97% identity with USP18. However, USP41 does not act as a deISGylase, which led us to perform a comparative analysis to decipher the basis for this difference, revealing molecular determinants of USP18’s specificity towards ISG15. We found that USP18 C-terminus, as well as a conserved Leucine at position 198, are essential for its enzymatic activity and likely act as functional surfaces based on AlphaFold predictions. Finally, we propose that USP41 antagonizes conjugation of the understudied ubiquitin-like protein FAT10 (HLA-<u>F</u> <u>a</u>djacent transcript <u>10</u>) from substrates in a catalytic-independent manner. Altogether, our results offer new insights into USP18’s specificity towards ISG15, while identifying USP41 as a negative regulator of FAT10 conjugation.

## INTRODUCTION

Post-translational modifications (PTMs) by members of the ubiquitin-like family of proteins is a fundamental aspect of cell signaling involved in many biological processes^1^. The founding member of this family, ubiquitin, is a small 8.5 kDa protein that gets conjugated to other proteins by using a cascade of enzymes termed E1, E2, and E3, which results in the covalent modification of a substrate with ubiquitin onto a lysine residue^2^. Formation of a polyubiquitin chain on a substrate often leads to its degradation by the proteasome, which is known as the ubiquitin-proteasome system (UPS)^2^. However, ubiquitination can also regulate protein-protein interactions, endocytosis, protein function or localization^1^. The complexity of ubiquitination is illustrated by the hundreds of enzymes involved in the conjugation process, as well as the different ways by which ubiquitin can be conjugated^3^, i.e. as a monomer (monoubiquitination), several monomers (multiubiquitination), and polyubiquitin chains linked through different linkages or with different topologies (homo- or heterotypic, as well as branched chains)^4^.

Ubiquitin-like proteins (Ubls) add another layer of complexity to this signaling apparatus^5^. These proteins share some sequence similarity to ubiquitin, as well as its three-dimensional structure^5^. Importantly, Ubls also get covalently attached to target proteins in a 3-step enzymatic cascade akin to ubiquitin, albeit with their own conjugation machinery^6^. The most well-known ubiquitin-like proteins include Nedd8^7^ (<u>n</u>eural precursor cell <u>e</u>xpressed, <u>d</u>evelopmentally <u>d</u>own-regulated <u>8</u>) and the SUMO^8^ (<u>s</u>mall <u>u</u>biquitin-like <u>mo</u>difier) family of proteins. However, the first Ubl that was shown to present the key feature of becoming covalently conjugated to intracellular proteins was ISG15 (interferon <u>s</u>timulated <u>g</u>ene <u>15</u>), in a process called ISGylation^9,10^. In contrast to ubiquitin, Nedd8 and SUMO, ISG15 structurally resembles two ubiquitin-like molecules that each share ∼30% sequence similarity to ubiquitin, and which are connected by a short linker, or hinge, region^11^. Moreover, while ubiquitin, Nedd8 and SUMO are all proteins that are constitutively expressed in cells, ISG15 is strongly induced by type I interferons^9^, such as IFN-α and IFN-β. Following interferon production, ISG15 gets translated as a 17 kDa precursor protein that is then processed into its mature 15 kDa form, to expose its carboxy-terminal LRLRGG motif necessary for its covalent attachment to target proteins^12^. The expression of the ISG15 conjugation machinery is also induced by interferon production, and this enzymatic cascade includes the E1 activating enzyme UBE1L/UBA7^13,14^, the E2 conjugating enzyme Ube2L6/UbcH8^15,16^, and HERC5 acting as the major E3 ligase^17,18^, even though a handful of other E3s have been described, such as ARIH1^19^, EFP^20^ (estrogen-responsive finger protein) and TRIM25^21^. In contrast to ubiquitin, ISGylation is not a proteolytic signal, and ISG15 gets conjugated as monomers on target proteins^22^. Moreover, ISG15 does not form chains, even though mixed ubiquitin-ISG15 chains have been described^23^. Instead, the main function of ISGylation is to contribute to the host response following bacterial^24^ and viral infections^13^. This is modulated in large part through co-translational attachment of ISG15 to viral capsid proteins, as well as host proteins, which ultimately inhibits virus assembly and/or replication^25^. Other agents causing ISG15 expression include lipopolysaccharide (LPS)^26^, retinoic acid^27^, as well as certain genotoxic stressors^28^, which all illustrate the role of ISG15 in triggering a host immune response to pathogenic stimuli.

Like ubiquitination, ISGylation is dynamic and can be reversed through the action of a specific cysteine protease called USP18^29^. While other proteases, including USP21^30^ and USP16^31^, have been reported to also deconjugate ISG15, it is believed that USP18 is the main deISGylating enzyme in vivo. Indeed, mouse USP18 (also named Ubp43) knock out cells display global accumulation of ISGylated proteins^26^ while USP18 overexpression leads to a dramatic decrease of protein ISGylation^32^. As with the rest of the ISG15 machinery, USP18 expression is also triggered by type I interferons^26^, and therefore acts as the major negative regulator of protein ISGylation. USP18 appears to show very little substrate specificity and will simply remove ISG15 from virtually any protein^29^. In addition to cleaving the covalent bond between ISG15 and a target protein, USP18 can also interacts non-covalently with ISG15^33^. Furthermore, USP18 expression can be stabilized by ISG15^33^. However, the basis for these modes of regulation is still unknown, and the molecular determinants responsible for the high specificity of USP18 towards ISG15 are not well understood. A recent study offered important insights into how USP18 engages ISG15 by solving the crystal structure of mUSP18/Ubp43 in complex with mouse ISG15^34^. This structure identified motifs called ISG15-binding box 1 (IBB-1) and 2 (IBB-2) on mUSP18/Ubp43, which recognize hydrophobic regions in the C-terminal ubiquitin-like domain of mISG15. But, since mUSP18/Ubp43 differs from human USP18 in its ability to bind ISG15^33^, additional mechanisms such as PTMs or binding partners might contribute to their Ubl specificity.

Using a mass-spectrometry-based approach, we identified USP41 as a USP18 binding protein. Interestingly, USP41 is a recently evolved paralog of USP18, present in humans, but not, for example, in mice, which possess the single USP18 ortholog, mUSP18/Ubp43. As paralog genes, human USP18 and USP41 share very high sequence identity (> 80%). Yet, remarkably, USP41 does not act as an ISG15 protease. A comparative analysis, which combined molecular biology, biochemistry, and in vitro enzymatic assays, allowed us to illuminate previously unknown molecular determinants of USP18 specificity towards ISG15. Our results identify the last 14 amino acids of USP18 on its C-terminus as being crucial for both its enzymatic activity and binding to ISG15. Importantly, the requirement of these last 14 amino acids is conserved across species since it is also critical for the enzymatic function of mUSP18/Ubp43. Moreover, we identify Leu198 on human USP18 as being important for ISG15 recognition. Finally, since USP41 is not a second ISG15 protease, we aimed at characterizing which Ubl it controls in vivo. Despite a few reports stating that it is a deubiquitinating enzyme (DUB)^35,36^, we found that USP41 can regulate the Ubl FAT10 (HLA-<u>F a</u>djacent transcript <u>10</u>), which presents similar features as ISG15. For instance, FAT10 structure is comprised of two ubiquitin-like molecules^37^, and its expression is induced by pro-inflammatory cytokines such as IFN-γ and tumor necrosis factor (TNF), while it is mostly restricted to cell types of the immune system^38^. We found that USP41 interacts non-covalently with FAT10 and that it can also downregulate the level of FAT10-conjugated proteins in cells. Until now, no gene product has been shown to downregulate FAT10 conjugation, and our results suggest that USP41 might fulfill this role in a catalytic-independent manner. Altogether, our data provide new insights into USP18’s specificity towards ISG15 which are conserved between human and mouse, while identifying USP41 as a negative regulator of FAT10. Our work highlights the power of performing a comparative analysis between paralog genes, like USP18 and USP41, to identify the biochemical basis for their differences and illuminate unique features of protein function. Importantly, USP18 and to a lesser extent USP41 have both been implicated in diseases. Thus, understanding their mechanisms of action might aid in the future development of inhibitors to impact disease progression.

## RESULTS

### Identification of the uncharacterized DUB USP41 as a USP18 interacting protein

The non-covalent interaction between USP18 and ISG15 has been well documented^33^, and interestingly, is unique to human proteins, since mUSP18/Ubp43 does not bind to murine ISG15 (Supplementary Figure 1A). Likewise, the stabilization of USP18 by ISG15 is also specific to humans, as previously reported^33^ (Supplementary Figure 1A, Input panel). Therefore, we reasoned that additional binding partners or post-translational modifications could impact USP18 binding to ISG15, in other ways than the ones already described for the interaction between mUSP18/Ubp43 and mISG15.

We initially aimed to identify PTMs on human USP18 when it is in complex with human ISG15. We co-transfected HEK-293T cells with a 6His-FLAG-tagged version of ISG15 (6HF-ISG15), along with an untagged version of USP18. The rationale for using an untagged form of USP18 is to allow for both long and short isoforms of USP18 to be produced in cells^39^ (Figure 1A, Input panel). We performed immunoprecipitation of ISG15 using FLAG-resin, after which the resulting immunoprecipitates were separated by SDS-PAGE and the gel was stained by Colloidal Coomassie Blue, allowing us to visualize that the two isoforms of USP18 readily co-precipitated with ISG15 (Figure 1A). The Coomassie-stained bands of both isoforms were cut out, digested, and analyzed by mass-spectrometry, and the corresponding areas in the USP18 only condition were used as negative controls. This analysis revealed that USP18 is not extensively modified with any PTMs while in complex with ISG15, as we only found Lys173 to be modified with a di-Gly (GG) peptide (Supplementary File 1). However, our analysis identified another member of the USP family, USP41, specifically in our ISG15 immunoprecipitates and not in the negative control (Figure 1B, Supplementary Figure 1B, and Supplementary Table 1 and 2, orange lines). USP41 shares significant sequence identity to USP18 and many identical tryptic peptides. However, we identified USP41 specific peptides by mass spectrometry (Figure 1B and Supplementary Figure 1B), confirming its endogenous expression in HEK293T. USP41 is a protein with a similar organization and molecular weight as USP18 (Figure 1C), suggesting its identification by mass-spectrometry is neither an experimental artifact nor a contamination. Moreover, since we detected USP41 following ISG15 immunoprecipitation, this observation suggested that USP41 might be another ISG15-interacting protein.

**Figure 1.**
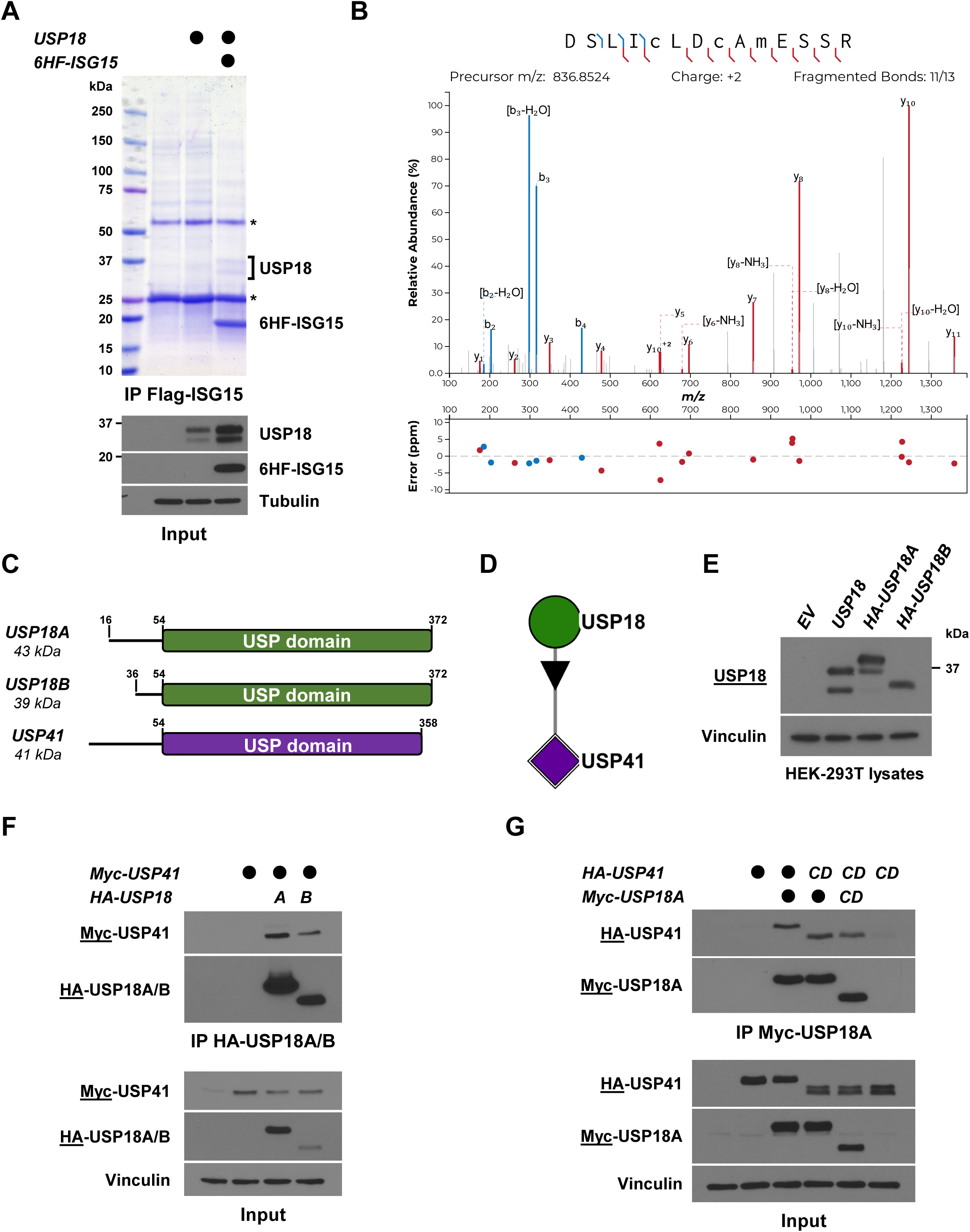
Identification of the uncharacterized DUB USP41 is a USP18 interacting protein. (a) Untagged USP18 was ectopically expressed in HEK-293T cells, either alone or with a 6His-FLAG-tagged ISG15 (6HF-ISG15). After 48h, precleared lysates were used to immunoprecipitate ISG15 on anti-FLAG beads. Immunoprecipitates were separated by SDS-PAGE and the gel was stained using Colloidal Coomassie Blue (CCB) staining, while inputs were analyzed by western blotting. The two USP18 isoforms that co-precipitated with ISG15 were excised, digested with trypsin, and sent for mass-spectrometry analysis. Asterisks indicate heavy and light chains of the FLAG antibody used for IP. (b) MS/MS spectrum of the doubly charged ion (m/z 836.8524) corresponding to USP41 tryptic peptide DSLICLDCAMESSR. Carbamidomethylation is present on C5 and C8 corresponding to C178 and C181 in USP41. Oxidation is present on M10 corresponding to M183 in UPS41. (c) Cartoon showing the organization of USP18A, USP18B and USP41. (d) Interaction network from the BioPlex Interactome database in HEK-293T cells, by querying USP18 as a gene. (e) HEK-293T cells were transfected with the indicated expression constructs and cell lysates were probed for USP18 using a monoclonal antibody raised against USP18 N-terminus (underlined). EV, empty vector. (f) Myc-USP41 and HA-USP18 (both isoforms) were ectopically expressed in HEK-293T cells. After 24h, precleared lysates of transfected cells were used to immunoprecipitate USP18A and B on anti-HA beads. IP and input samples were separated by SDS-PAGE and analyzed by western blotting. Immunoblotted antigen is underlined to the left of blots. (g) Same as in (f) except that USP18A was used, as well as the catalytic domains (CD) of both USP41 and USP18A (USP41 CD and USP18A CD, respectively).

To explore this possibility, we took advantage of publicly available protein-protein interaction databases. We used both the BioPlex (https://bioplex.hms.harvard.edu/) and BioGRID (https://thebiogrid.org/) databases and looked for interacting proteins after querying for ISG15. While both databases reported several identical ISG15-interacting proteins such as UBA7, the E1 for ISG15, or several members of the IFNA family of proteins (such as IFNA1, IFNA2, INFA8, IFNA10, IFNA14), neither database reported an interaction between ISG15 and USP41. Because the BioPlex includes interaction networks for nearly 15,000 proteins that were done in HEK-293T cells, the same cell line that we used for our IP-MS experiment, we used it to look at USP41 interacting proteins. Remarkably, by querying USP41 in the BioPlex, it is found as the only interactor of USP18 (Figure 1D). Moreover, by using the BioGRID and looking at USP18 binding proteins, USP41 is among the top interacting proteins, ranking alongside ISG15 or STAT2, which are well established interactors of USP18. Thus, this suggests that rather than interacting with ISG15, USP41 could be a USP18 binding protein.

As stated previously and as shown in Figure 1C, USP18 exists as two isoforms that we refer to as USP18A and USP18B, with starting sequences at positions 16 and 36, respectively. The reason the numbering of USP18 isoforms starts at these positions is because the USP18 reference sequence includes 15 amino acids on its N-terminus that are not part of the final gene products. This is because translation of human USP18 is not initiated at the predicted start codon at position 1 (AUG1). Instead, translation of USP18 isoforms is initiated both at the rare start codon CUG16 (USP18A) and at a second in-frame AUG36 (USP18B)^39^. Thus, we constructed HA-tagged versions of USP18A and B which reflect the correct expression of USP18 isoforms (Figure 1E and Supplementary Document 1 for protein sequences). To study their potential interaction, we conducted co-immunoprecipitation experiments in HEK-293T cells. We transfected a Myc-tagged version of USP41, alone or in combination with HA-tagged versions of USP18A or USP18B, and following immunoprecipitation of either USP18 isoform, we could observe co-immunoprecipitation of USP41 (Figure 1F). Importantly, this interaction was also observed by precipitating USP41 (Supplementary Figure 1C). Both USP18 and USP41 have a similar domain organization, where most of their sequences encompass their respective USP catalytic domain, which are preceded by a short N-terminus (Figure 1C). To map the regions that allow the USP18-USP41 interaction, we generated truncations of USP18 and USP41 lacking their respective N-termini, leaving only their USP catalytic domain (USP18 CD and USP41 CD, respectively). These truncated versions, as well as the full length (FL) versions of both enzymes, were transfected in HEK-293T cells and the interaction was monitored by co-immunoprecipitation. USP41 bound to USP18, and importantly, this binding occurs through their respective USP domains (Figure 1G and Supplementary Figure 1D). Taken together, these results identify the uncharacterized DUB USP41 as a novel interacting protein of USP18, whose binding depends on their respective catalytic domains.

### USP41 is a paralog of USP18 which does not react with ISG15

Compared to USP18, USP41 is a largely uncharacterized DUB. However, sequence analysis of USP41 revealed that it is a close paralog of USP18, as illustrated by their very high sequence identity (Figure 2A and Supplementary Document 1). This similarity led us to hypothesize that USP41 could be a second ISG15 protease. We first tested whether USP41 could remove ISG15 from endogenous proteins by using an overexpression system. This approach relies on transfecting HEK-293T cells with plasmids encoding 6HF-ISG15, together with its E1 (V5-UBA7) and E2 (V5-UbcH8). Co-expression of all 3 plasmids leads to conjugation of ISG15 to endogenous proteins, and this is not observed if the E1 is omitted, if the E2 is inactive (C86A), or if a non-conjugatable mutant of ISG15 (ΔGG) is used (Supplementary Figure 2A). This constitutes an ideal system to study deISGylation by proteases, and test whether USP41 could act as a deISGylating enzyme. Indeed, when using this system, we could observe that co-expression of USP18 led to the complete removal of ISG15 from endogenous proteins, and this is dependent on its catalytic cysteine (Figure 2B, compare USP18 WT and C64S lanes). In contrast, co-expression of USP41 had no effect on ISGylation in HEK-293T cells (Figure 2B). This is remarkable since the mouse ortholog of USP18, mUSP18/Ubp43, which has comparatively less sequence similarity to USP18 than USP41 (Supplementary Figure 2B), was able to deconjugate ISG15 from proteins as efficiently as human USP18 (Supplementary Figure 2C). To further assess the potential activity of USP41 towards ISG15, we performed two complementary in vitro assays. First, we used an ISG15 activity-based probe (ABP), ISG15-vinyl sulfone (ISG15-VS), whose C-terminal vinyl sulfone warhead will form a covalent bond with the active site cysteine of a protease, allowing us to monitor enzymatic activity. To do so, we first expressed HA-tagged versions of WT USP18 or USP41 in HEK-293T cells. The DUBs were then immunoprecipitated on HA resin and mixed with the ISG15-VS or with buffer as a control. As shown in Figure 2C, USP18 readily reacted with the ABP, as illustrated by its shift in size on SDS-PAGE. In striking contrast, USP41 did not react with the ISG15-VS ABP. As a complementary assay, we used the fluorogenic substrate ISG15-AMC which generates fluorescence following AMC cleavage by a protease. Immunoprecipitates of HA-tagged WT USP18, catalytically inactive USP18, or WT USP41 were mixed with the ISG15-AMC substrate. We observed an increase in fluorescence, indicative of protease-mediated cleavage of AMC from ISG15, only when the fluorogenic substrate was mixed with WT USP18, and not with USP41 or a catalytically inactive version of USP18 (Figure 2D and Supplementary Figure 6A for control of expression).

**Figure 2.**
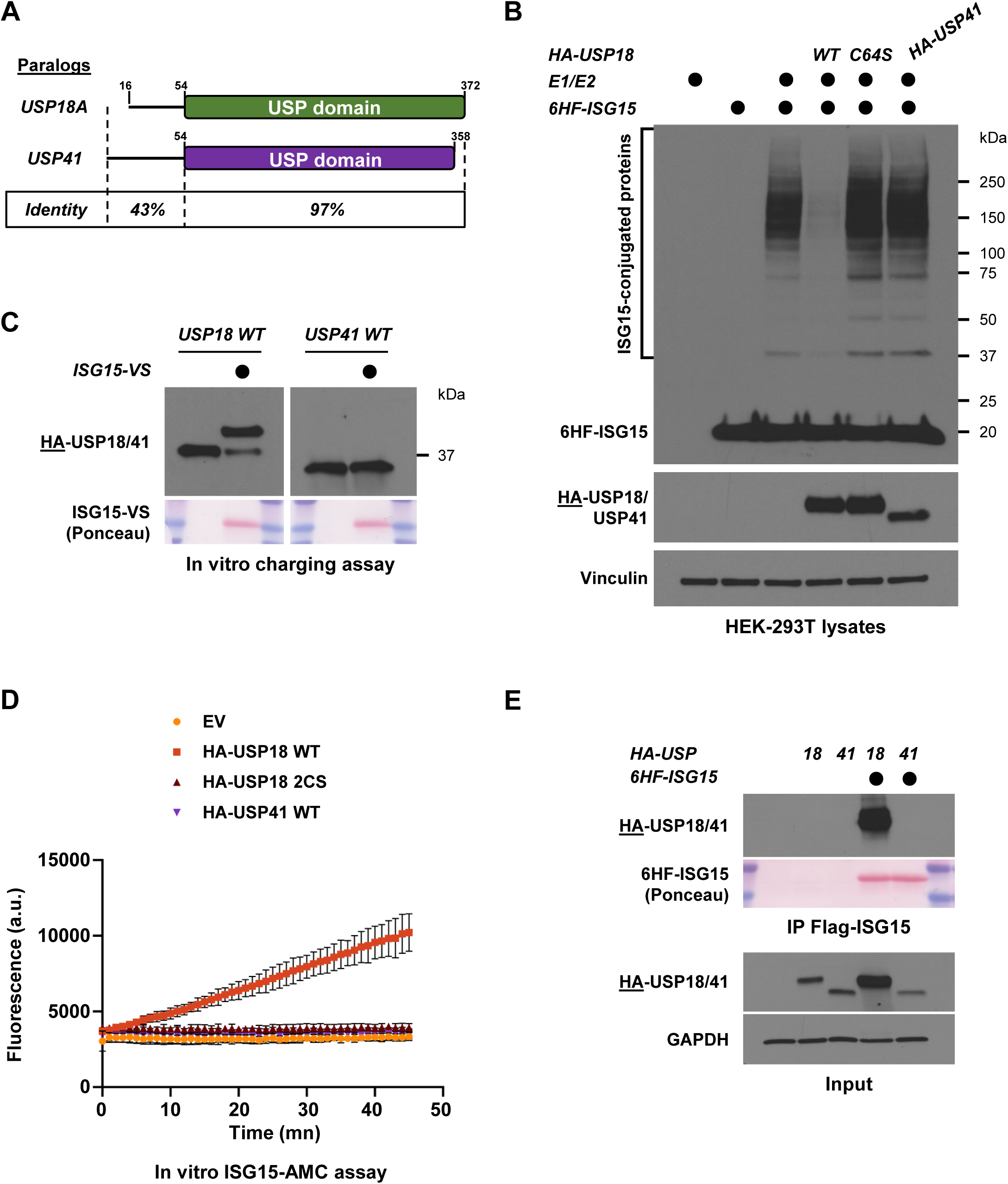
USP41 is a paralog of USP18 which does not react with ISG15. (a) Cartoon showing the sequence identity of USP18 and USP41 (b) Protein ISGylation in HEK-293T cells was reconstituted by ectopically expressing V5-UBA7 (E1), V5-UbcH8 (E2) and 6His-FLAG-ISG15 (6HF-ISG15). Where indicated, HA-USP18 WT, a catalytically inactive version (C64S), or HA-USP41, were co-transfected. After 24h, lysates of transfected cells were prepared then analyzed by SDS-PAGE and western blot. Immunoblotted antigen is underlined to the left of blots. (c) HA-USP18 and HA-USP41 were ectopically expressed in HEK-293T cells and after 24h, precleared lysates of transfected cells were used to immunoprecipitate USP18 or USP41 on anti-HA beads. Immunoprecipitates were mixed with reaction buffer containing an ISG15 activity-based probe (ISG15-VS) or not, and reaction products were analyzed by SDS-PAGE and western blot. (d) The indicated constructs were ectopically expressed in HEK-293T cells and after 24h, precleared lysates of transfected cells were used to immunoprecipitate USP18 WT, USP18 2CS or USP41 WT on anti-HA beads. Immunoprecipitates were mixed with reaction buffer containing the fluorogenic substrate ISG15-AMC, and fluorescence increase was monitored using a plate reader at the appropriate excitation and emission wavelengths. (e) HA-USP18 and HA-USP41 were ectopically expressed in HEK-293T cells, either alone or with 6His-FLAG-ISG15. After 24h, precleared lysates of transfected cells were used to immunoprecipitate ISG15 on anti-FLAG beads. Immunoblotted antigen is underlined to the left of blots.

Finally, we examined whether USP41 could bind ISG15, since USP18 non-covalently interacts with ISG15. To test this, HA-tagged versions of USP18 and USP41 were expressed, either alone or with 6HF-ISG15, and ISG15 was immunoprecipitated. As previously observed, we could readily detect USP18 interacting with ISG15, indicating that they interact non-covalently. In contrast, USP41 was not co-immunoprecipitated by ISG15 (Figure 2E). Moreover, whereas ISG15 co-expression led to a stabilization and increased abundance of exogenously expressed USP18 (Figure 2E, Input panel), it had no effect on USP41 protein levels. Thus, whereas USP18 binds, is stabilized by, and can deconjugate ISG15 in vivo, as has been extensively described, USP41 can neither react with, bind, nor be regulated by ISG15. Altogether, these data strongly suggest that despite being a paralog of USP18, USP41 does not act as a second ISG15 protease, even though their catalytic domains are 97% identical to each other.

### The C-terminus of USP18 is necessary for its enzymatic activity and binding to ISG15

The fact that USP41 cannot deconjugate ISG15, despite its high sequence similarity to USP18, led us to interrogate the basis for this enzymatic difference. When aligning the protein sequences of both enzymes, the most striking difference is the fact that the USP41 catalytic domain is shorter than USP18 by 14 amino acids on its C-terminus (Figure 2A). We therefore examined the impact of deleting these residues from either the FL version or the catalytic domain of USP18. To this end, we generated deletion constructs of USP18 (Figure 3A) and co-expressed each of them along with the ISG15 machinery in HEK-293T cells. As previously shown, USP18 can efficiently remove ISG15 from endogenous proteins, and this is dependent on its catalytic cysteine, since a C64S substitution impairs deISGylation in vivo (Figure 3B, compare FL to C64S). Strikingly, deleting the last 14 amino acids of USP18 completely abrogated its enzymatic activity (Figure 3B, compare FL to ΔC-term). Similarly, whereas the catalytic domain of USP18 retains the ability to deconjugate ISGylation in vivo, deleting the last 14 amino from the catalytic domain also led to impairment of USP18 activity (Figure 3B, compare CD to aa54-358). As a negative control, USP41 was also included and had no effect on ISG15-conjugated proteins. This indicates that the last 14 amino acids on the USP18 C-terminus (residues number 359-372) are crucial for its enzymatic activity.

**Figure 3.**
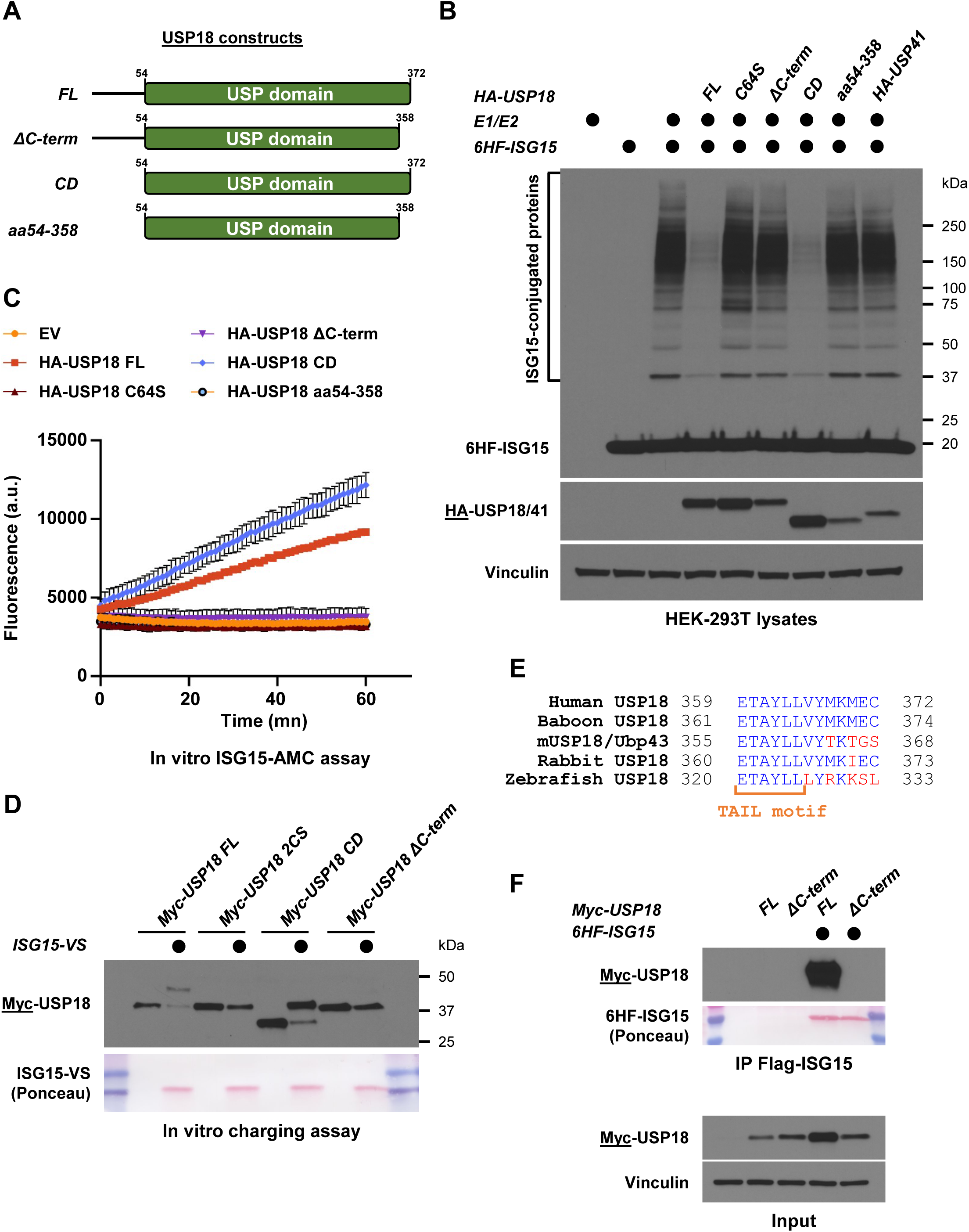
The C-terminus of USP18 is necessary for its enzymatic activity and binding to ISG15. (a) Cartoon showing the different truncation constructs of USP18 that were generated. The ΔC-term lacks the last 14 amino acids of the USP domain, the CD construct leaves only the catalytic domain, and the aa39-343 construct represents USP18 catalytic domain without the last 14 amino acids. FL, full length. (b) Protein ISGylation in HEK-293T cells was reconstituted by transfection of the ISG15 machinery (E1/E2/ISG15) and where indicated, HA-tagged USP18 constructs were co-transfected. After 24h, lysates of transfected cells were prepared then analyzed by SDS-PAGE and western blot. Immunoblotted antigen is underlined to the left of blots. (c) The indicated constructs were ectopically expressed in HEK-293T cells and after 48h, precleared lysates of transfected cells were used to immunoprecipitate HA-tagged USP18 variants on anti-HA beads. Immunoprecipitates were mixed with reaction buffer containing the fluorogenic substrate ISG15-AMC, and fluorescence increase was monitored as previously described. (d) The indicated Myc-tagged USP18 constructs were ectopically expressed in HEK-293T cells and after 48h, precleared lysates of transfected cells were used to immunoprecipitate USP18 on anti-Myc beads. Immunoprecipitates were mixed with reaction buffer containing ISG15-VS or not, and reaction products were analyzed by SDS-PAGE and western blot. (e) Sequence alignment of the C-termini of human USP18, baboon USP18, mUSP18/Ubp43, rabbit USP18 and zebrafish USP18. Red indicates residues that are divergent, while the TAIL motif is indicated in orange. (f) HA-USP18 FL or HA-USP18 ΔC-term were ectopically expressed in HEK-293T cells, either alone or with 6HF-ISG15. After 24h, precleared lysates of transfected cells were used to immunoprecipitate ISG15 on anti-FLAG beads. Immunoblotted antigen is underlined to the left of blots.

The above result led us to wonder if the importance of the USP18 C-terminus was conserved across species. We therefore turned our attention to the mouse ortholog of USP18, mUSP18/Ubp43. We co-expressed either FL, CD or ΔC-term versions of mUSP18/Ubp43 along with the human (Supplementary Figure 3A) or murine (Supplementary Figure 3B) ISG15 machinery. In both systems, deleting the last 14 amino acids of mUSP18/Ubp43 completely abrogated its enzymatic activity, to comparable levels as the catalytically inactive mutant C61S (Supplementary Figure 3A and B).

To further confirm the importance of the USP18 C-terminus for its enzymatic activity, we immunopurified our USP18 deletion constructs from HEK-293T cells and performed in vitro assays using the ISG15-AMC fluorogenic activity reporter and the ISG15-VS ABP. Both USP18 FL and USP18 CD were active towards ISG15-AMC, as evidenced by the increase in fluorescence over time (Figure 3C). Conversely, deleting the last 14 amino acids from either FL and CD USP18 constructs (ΔC-term or aa54-358, respectively) completely blocked activity towards ISG15-AMC (Figure 3C and Supplementary Figure 6B for control of expression). In a complementary experiment, the same constructs were used with the ISG15- VS ABP. Again, USP18 FL and USP18 CD reacted with the ABP, as shown by the formation of slower migrating forms of USP18 in the presence of the probe (Figure 3D). In contrast, the USP18 ΔC-term did not react with ISG15-VS (Figure 3D). Similarly, we also observed that mUSP18/Ubp43 FL could react with ISG15-VS ABP but not the ΔC-term mutant (Supplementary Figure 3C). Interestingly, a sequence alignment of human, baboon, mouse, rabbit, and zebrafish USP18 shows a high degree of conservation of USP18 C-terminus across species, further supporting a role for these residues in coordinating enzymatic function (Figure 3E). Moreover, we noticed that 6 evolutionary conserved residues form the sequence ETAYLL. Therefore, we name this patch the “TAIL motif” of USP18 (Figure 3E), and postulate that it is critical for USP18 activity towards ISG15, as suggested by its conservation among species and our biochemical analysis.

Finally, having established the importance of USP18 C-terminus in controlling its enzymatic activity, we then tested if it also affected USP18 ability to bind ISG15. To this end, HA-tagged versions of FL or ΔC-term USP18 were expressed, either alone or with 6HF-ISG15, and ISG15 was then immunoprecipitated on FLAG resin. In sharp contrast to FL USP18, the ΔC-term mutant was not co-immunoprecipitated by ISG15 (Figure 3F), indicating that the C-terminus of USP18 is also involved in binding to ISG15. By performing the reciprocal co-immunoprecipitation, we validated that ISG15 was not co-immunoprecipitated by USP18 ΔC-term (Supplementary Figure 3D). In addition, ISG15 co-expression led to increased abundance of the USP18 FL protein (Figure 3F, Input panel), and this ISG15-mediated stabilization of USP18 was not observed when its C-terminus was removed. Altogether, these results demonstrate that the C-terminus of USP18 is necessary for USP18 enzymatic activity and its ability to bind to ISG15. Moreover, we postulate that this is due to the presence of the TAIL motif, a stretch of 6 conserved amino acids that show a high degree of conservation across species (Figure 3E).

### Engineering USP41 into a deISGylase identifies Leu198 of USP18 as crucial for activity

Our finding that USP18 C-terminus is necessary for its enzymatic function prompted us to test if this would be sufficient to turn USP41 into an active ISG15 protease. To this end, we used USP18 and USP41 constructs expressing their catalytic domain only (Figure 4A), and we compared it to a chimeric construct of USP41 that is appended with the C-terminus of USP18 that we name USP41^+TAIL^ (Figure 4A). We expressed and purified these proteins from HEK-293T cells, then assessed their ability to react with the ISG15-VS ABP. Charging of USP18 was dependent on its C-terminus as previously observed (Supplementary Figure 4A). However, adding the TAIL motif to USP41 did not allow it to react with the ISG15-VS ABP (Supplementary Figure 4A). Thus, while the USP18 C-terminus is necessary for its enzymatic activity, it is not sufficient to turn USP41 catalytic domain into an ISG15 protease, despite sharing 97% identity with USP18, which suggests that other determinants are involved.

**Figure 4.**
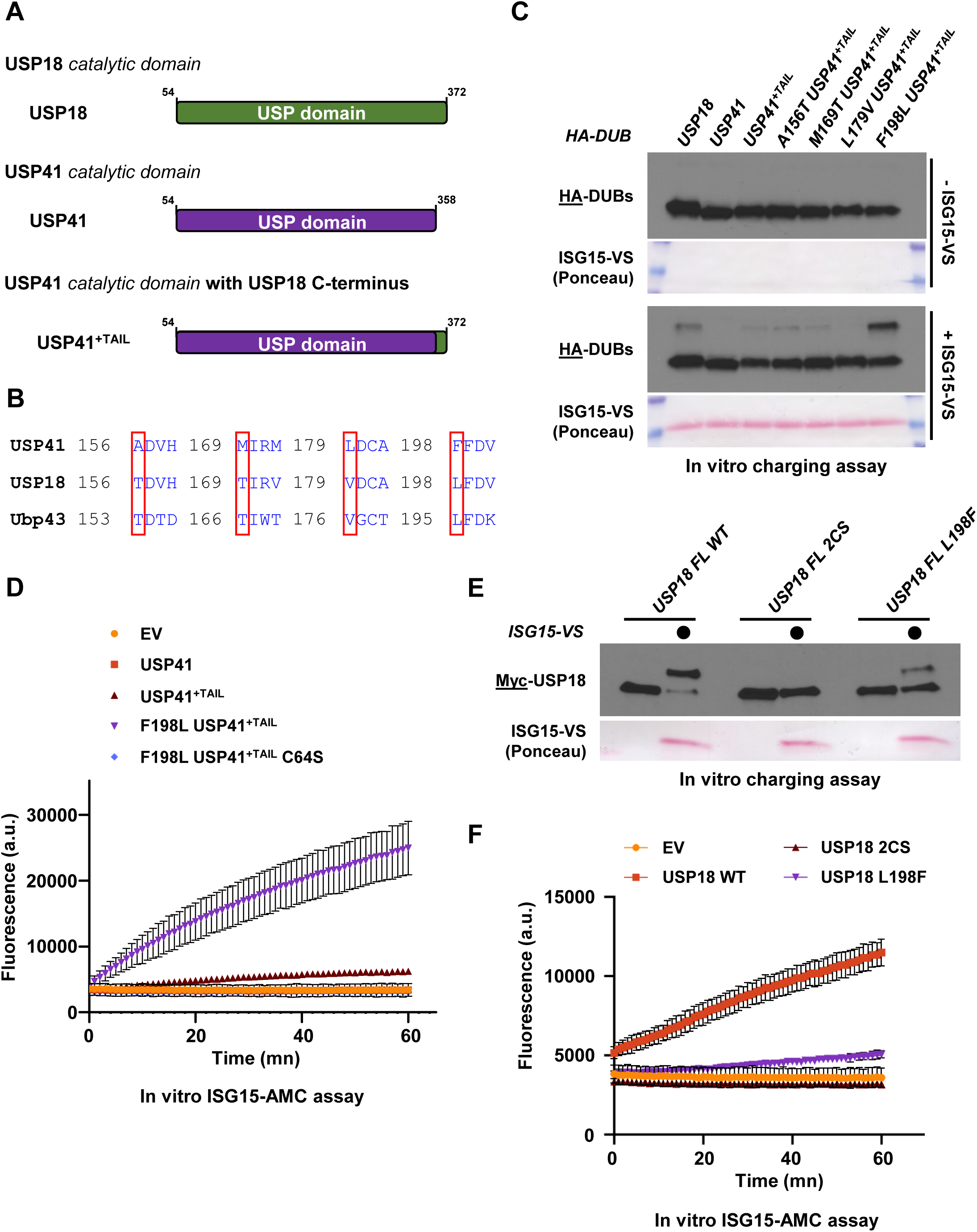
Engineering USP41 into a deISGylase identifies Leu198 of USP18 as crucial for activity. (a) Cartoon showing the different truncation constructs of USP18 and USP41 that were generated. USP41^+TAIL^ is a chimeric construct where the last 14 amino acids of USP18 were added to the USP domain of USP41. (b) Sequence alignment of Ubp43, USP18 and USP41. Red boxes highlight residues that are different between USP18 and USP41 but conserved between USP18 and mUSP18/Ubp43. (c) The indicated HA-tagged constructs were ectopically expressed in HEK-293T cells and after 24h, precleared lysates of transfected cells were used to immunoprecipitate these HA-DUBs on anti-HA beads. Immunoprecipitates were mixed with reaction buffer containing ISG15-VS or not, and reaction products were analyzed by SDS-PAGE and western blot. (d) Same as in (c), except that after immunoprecipitation, the indicated HA-tagged constructs were mixed with reaction buffer containing the fluorogenic substrate ISG15-AMC, and fluorescence increase was monitored as previously described. (e) The indicated Myc-tagged USP18 constructs were purified from HEK-293T cells on anti-Myc beads 48h after transfection. Myc-USP18 Immunoprecipitates were mixed with reaction buffer containing ISG15-VS or not, and reaction products were analyzed by SDS-PAGE and western blot. (f) The indicated Myc-tagged constructs were ectopically expressed in HEK-293T cells and after 24h, precleared lysates of transfected cells were used to immunoprecipitate Myc-tagged USP18 variants on anti-Myc beads. Immunoprecipitates were mixed with reaction buffer containing the fluorogenic substrate ISG15-AMC, and fluorescence increase was monitored as previously described.

To identify such determinants, we aligned USP18 and USP41 catalytic domains. Apart from their C-termini that differ in length, there are only nine amino acid differences, all within the first 150 residues of the USP catalytic domain. We reasoned that amino acids that are different between USP18 and USP41, but which are conserved between USP18 and mUSP18/Ubp43, would be important for deISGYlation. We therefore aligned USP18, USP41 and mUSP18/Ubp43, with the goal of finding such residues, and this revealed four candidates on USP41: Ala156, Met169, Leu179 and Phe198 (Figure 4B). We used the USP41^+TAIL^ chimeric construct to mutate these residues to the corresponding amino acid of USP18, and each new chimera was assayed using the ISG15-VS ABP. As previously observed, USP18 reacted with the probe, whereas USP41 or the chimera USP41^+TAIL^ did not (Figure 4C). Substitution of Ala156, Met169 or Leu179 to the corresponding USP18 residues (Thr156, Thr169 or Val179, respectively) did not turn USP41^+TAIL^ into a deISGylase. However, substitution of USP41 Phe198 to the corresponding Leucine of USP18 led to efficient charging of USP41^+TAIL^ with the ISG15-VS ABP (Figure 4C). Importantly, the newly engineered activity of USP41 towards ISG15 was dependent on its catalytic Cys64, which is conserved between USP18 and USP41, since mutating it to serine blocked charging with the ISG15-VS ABP (Supplementary Figure 4B). We sought to further validate the finding that the F198L substitution can indeed trigger enzymatic activity of the USP41^+TAIL^ chimera. First, we tested this newly engineered activity towards the ISG15-AMC fluorogenic substrate, as described above, and monitored the increase in fluorescence intensity as a readout for enzymatic activity. Consistently, AMC fluorescence increased only when the chimera F198L USP41^+TAIL^ was used, and again, this was blocked by mutating Cys64 into serine (Figure 4D and Supplementary Figure 6C for control of expression). Then, we co-expressed these constructs along with the ISG15 machinery in HEK-293T cells and monitored deISGylation of endogenous proteins. As previously observed, co-expression of USP18 led to the complete disappearance of ISG15 from proteins (Supplementary Figure 4C). Remarkably, the chimera F198L USP41^+TAIL^ was also able to cleave ISG15 from proteins in a similar manner as USP18, and this was dependent on Cys64 (Supplementary Figure 4C). Thus, the Phenylalanine at position 198 blocks USP41 activity towards ISG15.

To determine the importance of Leu198 for the activity of USP18 towards ISG15, we generated an L198F substitution and assessed the ability of this USP18 mutant to be charged with the ISG15-VS ABP or process the ISG15-AMC reporter. As observed previously, WT USP18 reacted with the ISG15-VS ABP, and this was fully blocked when the active site was mutated (Figure 4E, compare WT to 2CS). Remarkably, mutating Leu198 to Phenylalanine strongly reduced USP18’s ability to react with the ISG15-VS ABP (Figure 4E, compare WT to L198F). Consistently, the activity of USP18 L198F was severely impaired based on the ISG15-AMC fluorogenic reporter (Figure 4F and Supplementary Figure 6D, compare WT to L198F), further illustrating the importance of Leu198 for USP18 enzymatic activity. To establish the role of Leu198 in mediating USP18 activity, we also mutated this residue to phenylalanine on mUSP18/Ubp3 (L195F). Similar to what we observed with human USP18, a mutant version of mUSP18/Ubp43 harboring the L195F substitution significantly impaired the ability of Ubp43 to react with the ISG15-VS ABP (Supplementary Figure 4D), as well as the ISG15-AMC substrate (Supplementary Figure 4E and Supplementary Figure 6E for control of expression). Altogether, these findings demonstrate that on top of the TAIL motif, the protease activity of USP18 towards ISG15 is highly dependent on the presence of a conserved leucine residue at position 198 (human) or 195 (mouse).

### Structural analysis of USP18 Leu198 and TAIL motif using AlphaFold 3

Having established that the TAIL motif and Leu198 are important determinants of USP18 enzymatic activity, we next sought to put these findings in a structural context. As mentioned previously, the only available structural data for USP18 comes from X-ray crystallographic studies using the catalytic domain of mUSP18/Ubp43 in complex with mISG15^34^. Therefore, we took advantage of AlphaFold 3 to predict the structure^40^ of the catalytic domain of human USP18 in complex with human ISG15, where we highlighted Leu198 and the TAIL motif (Figure 5A). When overlayed with the crystal structure of mUSP18/Ubp43 in complex with mISG15 (PDB ID: 5CHV), the AlphaFold prediction of the human complex showed a good structural similarity with an overall rmsd of 0.60 Å. Strikingly, the TAIL motif is inserted directly into the catalytic domain, in between a structure made of several beta sheets that are located near the catalytic triad (Figure 5A), which highlights its importance for enzymatic function. On the other hand, Leu198 appears as an outlier, located on an unstructured stretch further away from the catalytic site (Figure 5A). However, a close-up view showed that it was located less than 10 angstroms away from ISG15 C-terminal part, leading us to wonder if Leu198 could be involved in mediating non-covalent interactions with ISG15 (Figure 5B).

**Figure 5.**
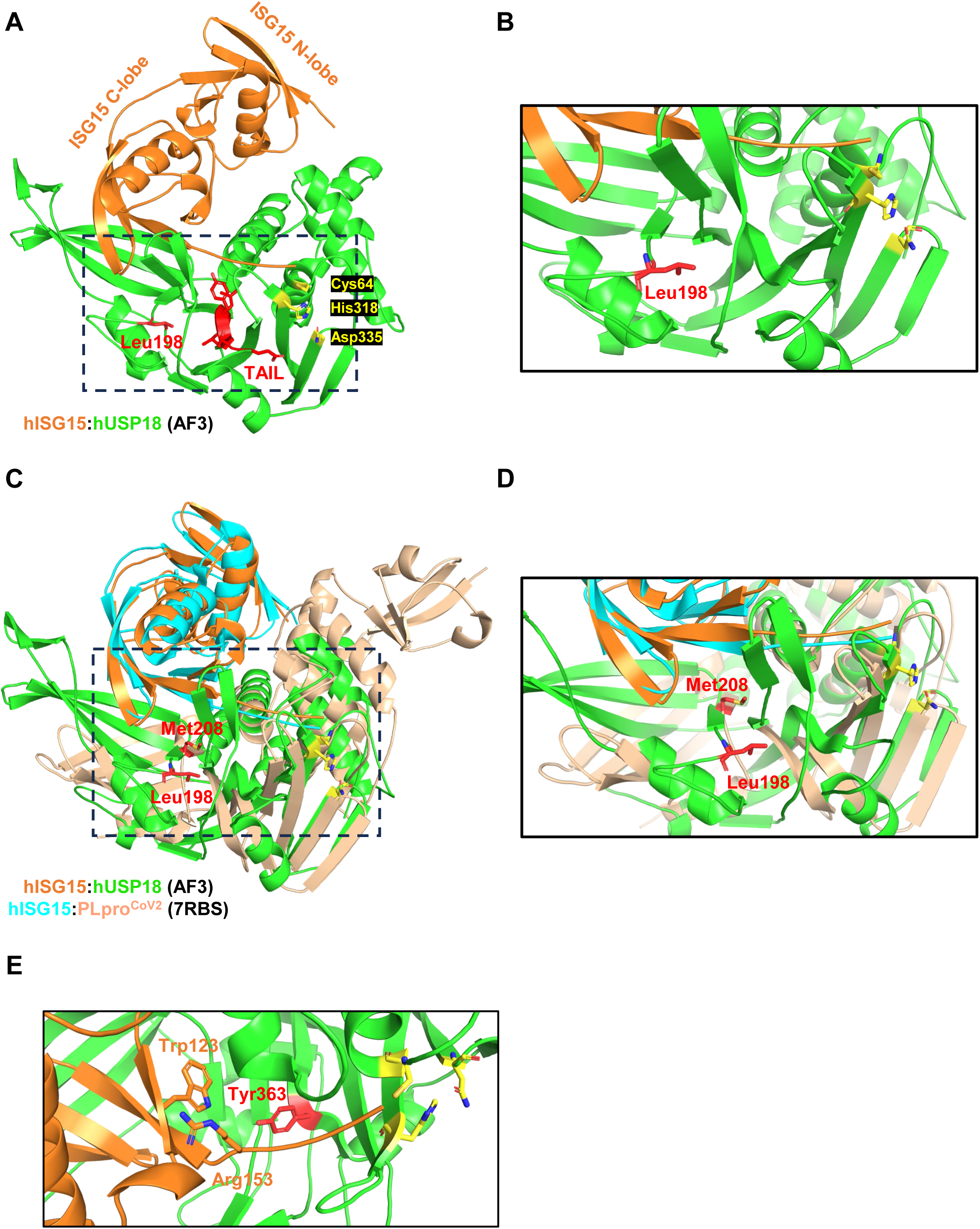
Structural analysis of USP18 Leu198 and TAIL motif using AlphaFold 3. (a) AlphaFold prediction of hUSP18 in complex with hISG15. Leu198 and the TAIL motif are highlighted in red, while the catalytic triad (Cys64, His318, Asn335) is in yellow. (b) Close up view of the dashed area in (a) showing Leu198 from hUSP18. (c) Overlay of hUSP18:hISG15 with the crystal structure of PLproCoV2:hISG15 (PDB identifier: 7RBS) by aligning the structures on ISG15. (d) Close up view of the dashed area in (c) showing Leu198 from hUSP18 and Met208 from PLproCoV2. (e) Close up view of the TAIL domain from hUSP18 with Trp123 and Arg153 of hISG15.

To explore this possibility and better understand the role of Leu198 in mediating USP18 activity, we turned our attention to the PLpro protease from SARS-CoV2, the virus responsible for COVID-19^41^. PLpro^CoV2^ is an open reading frame (ORF) located on the non-structural protein 3 (Nsp3) that has been shown to exert its function by acting as both a DUB and a deISGylase^42,43^. Interestingly, and despite low sequence identity, PLpro enzymes from various coronaviruses share common structural and catalytic properties with the family of Ubiquitin-specific proteases (USPs). Recently, the crystal structure of the PLpro^CoV2^ in complex with human ISG15 was solved^43^ (hISG15:PLpro^CoV2^, PDB ID: 7RBS), which gave us the opportunity to compare it to the AlphaFold structure of hISG15:hUSP18. We overlayed the two structural complexes by performing a pairwise alignment on the ISG15 molecules, which appeared nearly identical (Figure 5C, orange and cyan). When comparing the structures of both proteases, this analysis revealed that they differ significantly in their overall structure (Figure 5C). However, they also shared several structural features, such as a perfect alignment of their catalytic triads. Since key residues on PLpro^CoV2^ were previously identified to mediate non-covalent interactions with ISG15 (Tyr171, Glu167 and Met208)^43^, we looked for these and the corresponding ones on USP18. Interestingly, we found that Leu198 in USP18 is spatially located in the same area as Met208 in PLpro^CoV2^ (Figure 5D). This suggests that the presence of a small hydrophobic residue in that part of the protease structure is important for both human and viral mechanisms of deISGylation, where we assume that it helps coordinating ISG15 binding. Finally, the hISG15:PLpro^CoV2^ structure also identified three residues in ISG15 (Trp123, Pro130 and Arg153) that are important for interaction with PLpro^CoV2^. By looking at these amino acids on our hISG15:hUSP18 AlphaFold structure, we noticed that USP18 Tyr363, which is part of the TAIL motif, is located within 5 angstroms of both Trp123 and Arg153 of ISG15 (Figure 5E). Since these ISG15 residues were shown to be important for binding to the viral PLpro^CoV2^, this suggests that the TAIL motif might contribute to non-covalent interactions with ISG15. Altogether, by analyzing the AlphaFold structure of hISG15:hUSP18, we propose that both Leu198 and the TAIL motif of USP18 act as functional surfaces to recognize and bind conserved residues on ISG15.

### USP41 regulates conjugation of the Ubl FAT10 in an enzymatic-independent manner

Having established that USP41 function is likely to be independent of regulating ISG15 deconjugation, we sought to determine if it could regulate an alternative Ubl. To this end, we expressed and purified a Myc-tagged version of USP41 WT from HEK-293T cells, and mixed it with a panel of Ubl protein-ABPs harboring the vinyl sulfone warhead on their C-terminus (ISG15-VS, Ub-VS, HA-Ub-VS, Nedd8-VS or SUMO1-VS). USP41 was unreactive towards all ABP tested, which suggests that it is not regulating the conjugation of these ubiquitin-like proteins (Supplementary Figure 5A). However, this panel is not representative of all the ubiquitin-like molecules. Among other Ubls, we focused on FAT10 (HLA-<u>F a</u>djacent transcript <u>10</u>) since it is structurally similar to ISG15, made of two tandem Ub-like domains connected by a short linker region^37^. Because ISG15 is regulated by USP18, and USP41 is its closely related paralog with FAT10 being the closest ubiquitin-like protein to ISG15, we hypothesized that USP41 could regulate FAT10.

To test this, we used HEK-293 cells which have been shown to support FAT10 conjugation^44^. We first assessed binding between USP41 and FAT10 by co-immunoprecipitation experiment. We expressed HA-USP41 in HEK-293, either alone or with plasmids encoding 6HF-FAT10 or 6HF-ISG15. Following immunoprecipitation of both Ubl proteins on FLAG resin, we could observe interaction of USP41 with FAT10, but not ISG15 (Figure 6A). Conversely, when we performed the same experiment using USP18, we observed the opposite result where no interaction could be detected between USP18 and FAT10, and only between USP18 and ISG15 (Supplementary Figure 5B). The observation that USP41 readily interacts with FAT10 led us to test if USP41 also regulates deconjugation of FAT10 from endogenous proteins. The conjugation of FAT10 to substrates can be easily monitored by transiently transfecting HEK-293 cells with a 6His-FLAG-tagged FAT10 plasmid (6HF-FAT10). Indeed, following immunoprecipitation of FAT10 and FAT10-conjugated proteins on FLAG resin, we could observe a ladder of FAT10-conjugated proteins (Supplementary Figure 5C). As a control, ISG15 was also transfected and could not get conjugated to intracellular proteins (Supplementary Figure 5C). To test whether USP41 could act as a protease for FAT10 conjugation, we used the same approach and included conditions where either a WT or a mutant (2CS) USP41 harboring serine mutation on Cys64 and Cys65 were co-expressed with the 6HF-FAT10. Remarkably, co-expression of WT USP41 led to an almost complete disappearance of FAT10-conjugated proteins and surprisingly, this downregulation was not dependent on USP41 catalytic activity (Figure 6B, compare WT to 2CS). Importantly, we could show that both WT and 2CS versions of USP41 were co-immunoprecipitated by FAT10 (Figure 6B, IP panel), further supporting the notion that this downregulation happens independently of the enzymatic status of USP41.

**Figure 6.**
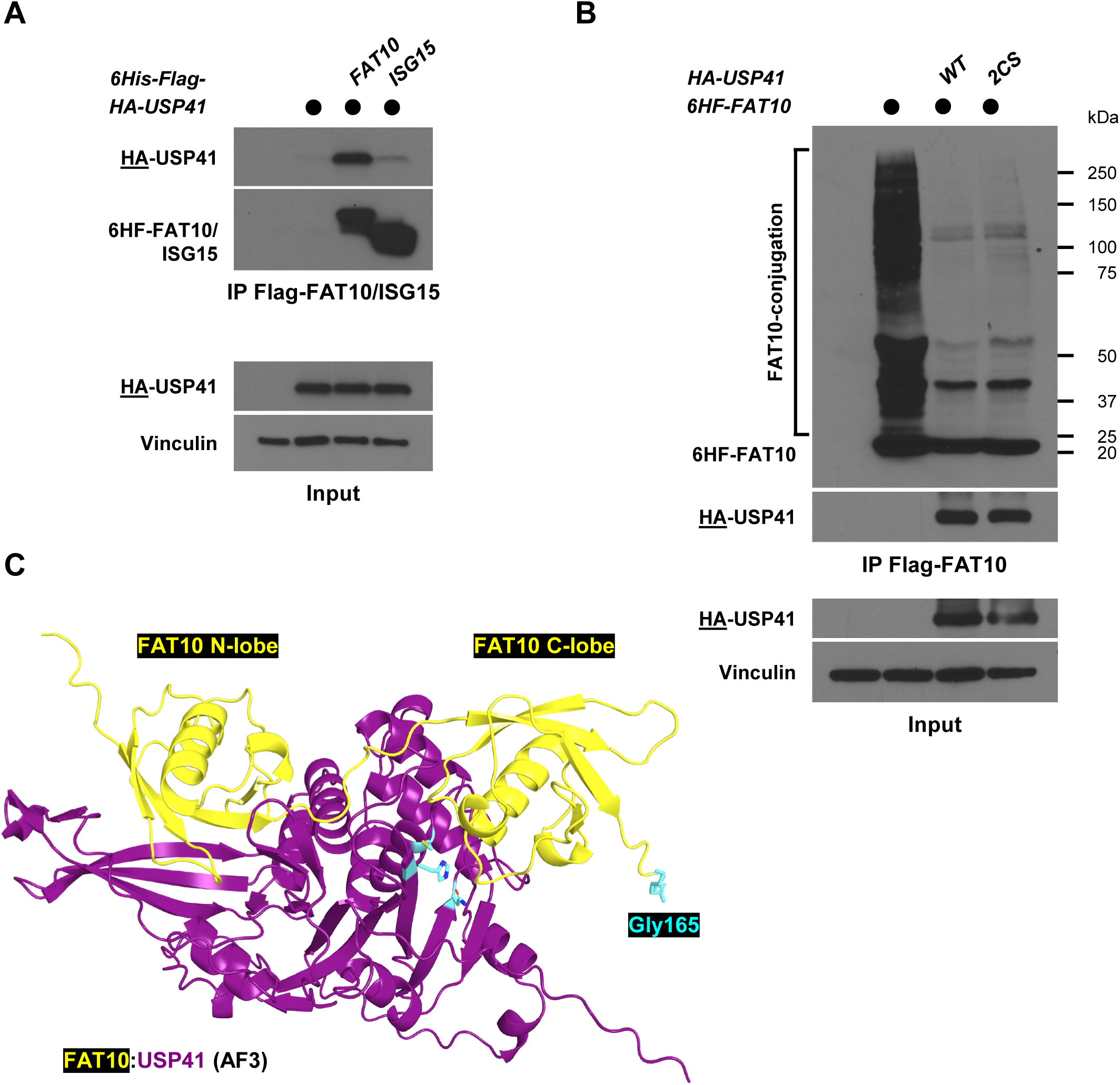
USP41 regulates conjugation of the Ubl FAT10 in an enzymatic-independent manner. (a) HA-USP41 was ectopically expressed in HEK-293 cells, either alone or with 6HF-FAT10 or 6HF-ISG15. After 24h, precleared lysates of transfected cells were used to immunoprecipitate FAT10 and ISG15 on anti-FLAG beads. Immunoblotted antigen is underlined to the left of blots. (b) 6HF-FAT10 was ectopically expressed in HEK-293 cells and where indicated, HA-tagged WT or a catalytically inactive version (2CS) of USP41 were also co-transfected. After 48h, precleared lysates were used to analyze FAT10 conjugation by immunoprecipitating FAT10-conjugated proteins on anti-Flag beads. IP and Input samples were analyzed by SDS-PAGE and western blot. (c) AlphaFold prediction of USP41 in complex with FAT10. The catalytic triad (Cys64, His318, Asn335) of USP41, as well as FAT10 Gly165 necessary for conjugation, are colored in cyan.

The observation that USP41 can regulate FAT10 conjugation in an enzymatic-independent manner led us to wonder if this feature could be explained at the structural level. Again, we took advantage of AlphaFold to generate the structure of USP41 bound to FAT10, where we highlighted the catalytic triad of the DUB as well as FAT10 Gly165, which gets conjugated to substrates (Figure 6C). Remarkably, the AlphaFold prediction of the FAT10:USP41 complex revealed that Gly165 is located far away from the predicted catalytic triad of USP41 (Figure 6C, cyan). This is in sharp contrast with the hISG15:hUSP18 complex, where ISG15 C-terminus is docked on USP18 active cysteine (Figure 5A and B), as it has been extensively described for cysteine-based DUBs. Finally, we asked how the two complexes differ, so we overlayed FAT10:USP41 with hISG15:USP18 by aligning them on the DUBs. Overall, the structure of both USP18 and USP41 was highly similar, showing a perfect alignment of their catalytic domains (Supplementary Figure 5D). However, we noticed that FAT10 N-terminal ubiquitin-like domain overlayed with ISG15 C-terminal ubiquitin-like domain, thus placing the FAT10 C-terminus away from USP41 catalytic triad, consistent with a catalytic independent mechanism of regulation (Figure 6B and 6C). Overall, these observations identify USP41 as a negative regulator of FAT10 and suggest that it antagonizes FAT10 conjugation in a catalytic-independent manner.

## DISCUSSION

USP18 is part of the USP family of DUBs, which is the largest sub-family of ubiquitin cysteine proteases^45^. Despite its name, and some reports describing that it regulates ubiquitination of a few substrates^46,47^, the most documented function of USP18 is to cleave the Ubl ISG15 from intracellular proteins^32^. Moreover, human USP18 can non-covalently bind to ISG15 with very high affinity^33^, yet the molecular basis for this remarkable specificity of USP18 towards ISG15 has remained elusive.

To better understand the basis for USP18’s extremely high affinity and specificity towards ISG15, we initially aimed at finding PTMs on USP18 when it is in complex with ISG15, by performing mass spectrometry analysis on USP18 after precipitating ISG15 from cells. Instead, we identified USP41 as a USP18 interacting protein, a finding that was corroborated by protein-protein interaction databases and additional coIP experiments (Figure 1). USP41 is a largely uncharacterized DUB and also a close paralog of USP18, illustrated by their high sequence homology (Figure 2A) and location on the same chromosome, indicative of a gene duplication. Despite being related to one another, USP41 does not act as an ISG15 protease like USP18, and at this point the function of their interaction remains to be elucidated. However, the very high sequence similarity between the catalytic domains of the two gene products (>97%) offered us the unique opportunity to decipher the basis for such different enzymatic activity, with the goal of characterizing new molecular determinants of USP18’s specificity for ISG15. By combining molecular and biochemical approaches, we identified the last 14 amino acids on USP18 C-terminus as being critical for its function. Indeed, a major finding of our study is the fact that USP18 C-terminus is necessary for enzymatic activity, since its deletion renders both USP18 or mUSP18/Ubp43 completely inactive. Sequence alignment revealed the presence of a stretch of 6 amino acids that are conserved across species and form the sequence ETAYLL, which we name the TAIL motif (Figure 3E).

The importance of the TAIL motif is particularly interesting when considering the various mechanisms of action of USP18. On top of its role in deconjugating ISG15 from proteins in an enzymatic-dependent manner, mUSP18/Ubp43 can also negatively regulate interferon signaling in an enzymatic-independent manner^48^. Through binding to the second chain of the interferon α/β receptor 2 (IFNAR2) complex, USP18 competes with Janus kinase 1 (JAK1), leading to downregulation of the JAK-STAT pathway and negative regulation of IFN-1 signaling^48^. Interestingly, it was shown that the C-terminus of USP18 mediates the interaction with IFNAR2, and the authors suggested that the residues within the last 50 amino acids of USP18 might be critical for this interaction. Remarkably, a few years after this report came out, a study by Richer & al. reported that a missense mutation in the TAIL motif of mUSP18/Ubp43 severely impacted mice susceptibility to *Salmonella typhimurium* infection^49^. By performing an in vivo screen using N-Ethyl-N-Nitrosourea (ENU) mutagenesis, the authors identified a loss-of-function mutation within the TAIL motif of mUSP18/Ubp43, where replacing Leu361 (Leu364 in human USP18) to phenylalanine conferred increased susceptibility to *S. typhimurium*^49^. Mechanistically, this phenotype was attributed to enhanced IFN-1 signaling, resulting in STAT1 hyperactivation and subsequent impairment of STAT4 phosphorylation and IFN-γ production. It is thus tempting to speculate that while the regulation of IFNAR2 signaling and ISG15 conjugation by USP18 are enzyme-independent and enzyme-dependent processes, respectively, they could still be coordinated by the same mechanism which involves the TAIL motif. However, additional studies will be needed to address the exact contribution of USP18 C-terminus, particularly the TAIL motif, in mediating the different mechanisms of action of USP18.

Even though the TAIL motif appears necessary for USP18 enzymatic activity, it is not sufficient to turn USP41 catalytic domain into an ISG15 protease, since our chimeric construct USP41^+TAIL^ was still inactive towards ISG15. By performing amino acids swapping (Figure 4B), we demonstrate using the same USP41^+TAIL^ chimera that Leu198 of USP18 is required for enzymatic activity, as this is sufficient to turn the inactive USP41^+TAIL^ construct into an active ISG15 protease (Figure 4 and Supplementary Figure 4).

Accordingly, mutation of Leu198 (Leu195 in mouse) to phenylalanine strongly reduced enzymatic activity of both USP18 and mUSP18/Ubp43 (Figure 4E and 4F), highlighting the importance of that residue for proper enzymatic function. To put these findings in a structural context, we took advantage of AlphaFold 3 to generate the structure^40^ of human USP18 catalytic domain in complex with ISG15 and compared it to the crystal structure of the COVID19 protease PLpro^CoV2^, which has been shown to act as a deISGylase. Interestingly, Leu198 of USP18 appears to be in the same space as Met208 of PLpro^CoV2^, which was found to be one of the key residues of the protease to interact with ISG15. Moreover, Trp123 and Arg153 on ISG15 were identified to mediate interaction with PLpro^CoV2^. Remarkably, our AlphaFold prediction of the hISG15:hUSP18 complex showed that Tyr363 of USP18, which is part of the TAIL motif, could be involved in interacting with these residues on ISG15 given its close proximity with Trp123 and Arg153.

Interestingly, these features were not reported when the crystal structure of the catalytic domain of mUSP18/Ubp43 in complex with mouse ISG15 was solved^34,50^. The main finding of this study explaining USP18’s selectivity was the identification of a hydrophobic domain called ISG15-binding box 1 (IBB1). IBB1 forms a contiguous patch made of 4 residues (Ala138, Leu142, Ser192 and His251) that appears crucial for interacting with a hydrophobic region unique to ISG15 located on its C-terminus Ubl domain. Mutation of IBB1 can effectively impair USP18 enzymatic activity in vitro, making it clear that these amino acids participate in ISG15 recognition. However, IBB1 is present on the catalytic domain of USP41 and yet, our chimera USP41^+TAIL^ did not show any enzymatic activity (Supplementary Figure 4A). Therefore, this clearly suggests that the presence of IBB1 alone is not sufficient to explain USP18’s activity towards ISG15, and our data provide new insights into USP18 mechanism of action.

It is important to emphasize that our findings that both Leu198 and the TAIL motif are important for USP18 enzymatic activity were facilitated by performing a comparative analysis with its paralog USP41. We found that despite their similarity, USP41 displays no apparent reactivity towards ISG15. Overall, USP41 is a vastly uncharacterized protein whose biological function remains largely unknown, but it is predicted to exhibit deubiquitinating activity based on the presence of the typical catalytic triad domain including Cys, His and Asn residues common to all cysteine DUBs. It has been reported that USP41 promotes proliferation of lung cancer cells^51^, as well as in other cancerous cell lines, through regulating the deubiquitination of the receptor for activated C kinase 1 (RACK1)^35^ and the epithelial-mesenchymal transition transcription factor (EMT-TF) Snail^36^. However, we failed to observe any reactivity of USP41 in vitro with a panel of activity-based probes, including ubiquitin (Figure 5A). Instead, we show that it binds and regulates FAT10, another Ubl similar to ISG15 in the sense that it looks like a ubiquitin dimer, and FAT10 is involved in response to type II Interferon, such as IFN-γ^38^. Interestingly, we found that the regulation of FAT10 conjugation by USP41 is independent of its predicted active cysteine (Figure 5D). By using AlphaFold 3 to generate an FAT10:USP41 complex, we observed that Gly165 of FAT10, which gets conjugated to proteins, is located far away from USP41 catalytic triad (Figure 6C and Supplementary Figure 6D). While surprising, this observation could be explained by the fact that several DUBs perform non-catalytic, or so-called “moonlighting” functions^52^, which extend beyond their ability to hydrolyze ubiquitin and Ubls. As described above, USP18 regulates IFN-1 signaling in a moonlighting manner, by virtue of its binding to INFAR2^48^. Another prominent example includes OTUB1 from the OTU (<u>O</u>varian <u>Tu</u>mor) family of DUBs, which can negatively regulate ubiquitination in an enzymatic-independent mechanisms^53^. Through binding to a subset of ubiquitin-charged E2s, OTUB1 prevents ubiquitin transfer by trapping the ubiquitin-charged E2. Based on our data and the AlphaFold prediction, it is tempting to speculate that USP41 might be sequestering FAT10 away from its conjugation machinery, leading to decreased levels of FAT10 conjugation, but this will require additional investigation.

Finally, it is notable that over the course of our study, USP41 went from being annotated as a protein-coding gene in databases to now being a pseudogene. The notion that USP41 could be a pseudogene was suggested in a study done by the Pellegrini lab, in which the authors argued that the USP41 gene lacks both 5’ and 3’ UTRs, on top of not finding evidence of USP41 transcripts in human tissues^54^. However, an early study which performed whole transcriptomic profiling by RNA-sequencing (RNA-seq) of macrophages stimulated with lipopolysaccharides (LPS) found USP41 to be one of the most upregulated genes following LPS activation^55^. Furthermore, we were able to detect protein peptides specific to USP41 in our mass-spectrometry experiment (Figure 1B and Supplementary Figure 1B), and the BioPlex database further supports the existence of USP41 in HEK-293T cells. Therefore, the mechanisms by which USP41 gets transcribed and translated might require further investigation. Another possibility explaining the scarcity of evidence regarding the existence of USP41 could lie in its expression pattern. Since it appears that USP41 regulates FAT10, whose expression is itself limited to a few cell types of the immune system^38^, we hypothesize that USP41 expression might also be limited to certain tissues, which has been observed in an early study that cloned and analyzed 22 USPs including USP41^56^. This will represent the basis for future work aiming at delineating the function, mechanisms of action, and distribution of USP41. Nevertheless, our results clearly suggest that USP41 acts as a negative regulator of FAT10 conjugation, and its high sequence homology with USP18 offered us a great opportunity to get new insights into the molecular determinants of USP18 specificity towards ISG15.

## MATERIALS AND METHODS

### Mammalian cell culture, transfection, cell lysis, antibodies, and reagents

HEK-293T cells were obtained from ATCC and grown in DMEM complete medium (Gibco, Cat# 11-965-092) supplemented with 10% fetal bovine serum (Atlanta Biologicals). HEK-293 cells were a kind gift from Dr Nancy Raab-Traub (UNC Chapel Hill, NC USA) and were grown in MEM (Gibco, Cat# 11-095-080) supplemented with 10% fetal bovine serum (Atlanta Biologicals). All DNA transfection experiments performed in HEK-293T and HEK-293 were done using PolyJet In Vitro DNA Transfection Reagent (SignaGen Laboratories) according to the manufacturer’s instructions, and subsequently cultured for 24 to 48h prior to analysis. Samples for protein analysis by immunoblot were lysed in NETN buffer [20 mM Tris-Cl (pH 8.0), 100 mM NaCl, 0.5 mM EDTA and 0.5 % (v/v) Nonidet P-40 (NP-40)] supplemented with 2 µg/ml pepstatin, 2 µg/ml apoprotinin, 10 µg/ml leupeptin, 1 mM AEBSF [4-(2 Aminoethyl) benzenesulfonyl fluoride], and 1 mM Na3VO4. Lysis was performed on ice for ∼10 minutes with occasional vortexing, lysates were then spun down at 14,000 rpm for 10 mn before determining protein concentration using Bradford reagent (Bio-Rad).

Standard immunoblotting procedures were followed. A list of reagents and antibodies that were used in this study, including the concentrations at which they were used, is available in Supplementary Table 3. All antibodies were diluted in 5% nonfat dried milk [diluted in Tris buffered saline, 0.05% tween-20 (TBST)], incubated for 1 hour at room temperature or overnight at 4°C, and detected using HRP conjugated secondary antibodies (Jackson Immuno Research Laboratories Inc; 1:5000 dilutions).

### ISG15 immuno-precipitation and mass spectrometry analysis

#### Sample preparation

For mapping PTMs on USP18 by mass spectrometry after ISG15 immuno-precipitation, 3 sets of 10x 10 cm dishes of HEK-293T cells were transfected with a total of 5 ug per plate (2.5 µg of untagged USP18, either alone or with 2.5 µg of 6HF-ISG15) and using PolyJet reagent according to the manufacturer’s instructions. Cells were grown for 48 hours, washed with PBS, and harvested in PBS then centrifuged at 1,500g for 5 mn. Cell pellets were lysed in NETN lysis buffer supplemented with protease and phosphatase inhibitors, as described above. Lysates were also snap frozen twice in liquid nitrogen, and then lysates were clarified by centrifugation at 14,000 rpm for 10 minutes at 4°C. Protein concentration was normalized using Bradford reagent. Samples were immunoprecipitated using EZview Red anti-FLAG M2 affinity gel (Millipore Sigma) for 2 hours at 4°C on a rotary shaker. Following IP, samples were washed 3x using NETN buffer followed by 3 washes using dPBS (Gibco, Cat# 14190144). Samples were resuspended in 2X Laemmli buffer, separated by SDS-PAGE on a pre-cast 4–15% Criterion™ Protein Gel (Bio-Rad Laboratories, Cat#5671084), and the gel was stained using QC Colloidal Coomassie Stain (Bio-Rad) according to the manufacturer’s instructions. The bands corresponding to UPS18 that co-precipitated with ISG were excised into 1 mm cubes, and destained for 20 minutes. Gel bands were washed with ACN, reduced with 10 mM DTT, alkylated with 100 mM iodoacetamide, and digested in-gel with 0.5 ug/uL trypsin overnight at 37°C. Peptides were extracted with ACN, desalted with Pierce C18 spin columns (ThermoFischer Scientific, Catalog # 69705) and dried via lyophilization. Peptide samples were stored at -80°C until further analysis.

#### LC-MS/MS Analysis

The peptide samples were analyzed by LC/MS/MS using an Easy nLC 1200 coupled to a QExactive HF mass spectrometer (ThermoFischer Scientific). Samples were injected onto an EasySpray column (75 μm id × 25 cm, 2.0 μm particle size) and separated over a 45 min method. The gradient for separation consisted of 5–45% mobile phase B at a 250 nl/min flow rate, where mobile phase A was 0.1% formic acid in water and mobile phase B consisted of 0.1% formic acid in 80% ACN. The QExactive HF was operated in data-dependent mode where the 15 most intense precursors were selected for subsequent fragmentation. Resolution for the precursor scan (m/z 350–1700) was set to 120,000 with a target value of 3 × 106 ions. MS/MS scans resolution was set to 15,000 with a target value of 1 × 105 ions. The normalized collision energy was set to 27% for HCD. Dynamic exclusion was set to 30 s, peptide match was set to preferred, and precursors with unknown charge or a charge state of 1 and ≥ 6 were excluded.

#### Data Analysis

Raw data files were processed using Sequest HT within Proteome Discoverer version 2.5 (ThermoFischer Scientific). Data were searched against a reviewed Uniprot human database (downloaded in February 2020), appended with a common contaminants database (MaxQuant, 245 sequences). The following parameters were used to identify tryptic peptides for protein identification: 10 ppm precursor ion mass tolerance; 0.02 Da product ion mass tolerance; up to two missed trypsin cleavage sites; (C) carbamidomethylation was set as a fixed modification; (M) oxidation, (K) Acetylation, (K) Ubiquitination, and (S/T/Y) phosphorylation were set as variable modifications. Peptide false discovery rates (FDR) were calculated by the Percolator node using a decoy database search and data were filtered using a 1% FDR cutoff. MS/MS spectra corresponding to unique peptides mapping to UPS41 (identified as the co-precipitated protein) were annotated using IPSA^57^.

### Molecular biology

USP41 and FAT10 were synthesized by Integrated DNA Technologies (Coralville, IA, USA) and their sequences can be found in the Supplementary Document 1 (gBlocks and protein sequences). FLAG-UbcM8, His6-mISG15 and HA-mUbp43 were gifts from Dr Dong-Er Zhang (Addgene plasmid #12440, #12445 and #12454, respectively), while FLAG-HA-USP18 was a gift from Dr Wade Harper (Addgene plasmid #22572). The ISG15 E1 (UBA7) and E2 (UbcH8) enzymes were kind gifts of Dr Robert Krug (University of Texas at Austin, TX USA). These plasmids served as templates to generate all the constructs used in this study, which were subcloned using restriction enzymes-based cloning into pcDNA3.1(+), a kind gift from Dr David Allison (UNC Chapel Hill, NC USA). PCR amplification of DNA was performed using Q5 High Fidelity DNA polymerase purchased from New England Biolabs. All PCR primers contained the appropriate restriction sites, followed by a kozak sequence, an appropriate N-terminal HA, Myc or V5 tag, and a di-glycine linker (see Supplementary Table 3 for details about plasmids and primers). All primers for site-directed mutagenesis were designed by using the NEBaseChanger website from New England Biolabs (https://nebasechangerv1.neb.com/) and following their protocol. To engineer the chimeric USP41^+TAIL^ constructs, we took advantage of the EcoRI restriction site present at position 787 of both genes. First, we PCR amplified a USP18 fragment, which started at the EcoRI site and covered the rest of USP18 sequence, while adding an XhoI site after the stop codon. Then, USP41 constructs were digested with EcoRI and XhoI, and ligation was done with the USP18 fragment. All constructs were verified by sequencing.

### Immuno-precipitation

Immuno-precipitation experiments (IP) were performed essentially as described previously^58^. Briefly, HEK-293T cells were transfected in 60-mm dishes with a total of 2.5 µg of DNA constructs for 24 to 48h. Cells were washed and scrapped in PBS then lysed for 10 min at 4°C with occasional vortexing in NETN buffer supplemented with protease inhibitors. Cell debris was removed by centrifugation at 14,000 rpm for 10 min at 4°C. Inputs were prepared with the clarified lysate, then anti-FLAG, anti-HA or anti-Myc beads (20 µl per IP, all from Millipore Sigma) pre-washed with lysis buffer were mixed with the remaining lysate and incubated on a rotary shaker for 1h to 1h30 at 4°C. Beads were washed with lysis buffer four times, resuspended in 25 ul of 2X sample buffer (60 mM Tris pH 6.8, 2% SDS, 10% glycerol, 100 mM DTT and 0.1% Bromophenol Blue) then incubated at 95°C for 5 minutes before SDS-PAGE and western blot analysis.

### Overexpression system to study ISGylation in HEK-293T cells

Generation of broad protein ISGylation in cells was performed by using an overexpression approach in HEK-293T cells based on a previously described protocol^59^, with minor changes. Briefly, HEK-293T cells were first seeded in 60 mm dishes, at approximately 3 million cells per dish. The day after, cells were transfected with a mix of 0.375 µg of V5-UBA7 (E1), 0.375 µg of V5-UbcH8 (E2) and 0.75 µg of 6HF-ISG15. Where indicated, 0.75 µg of the indicated DUBs constructs was also included, otherwise empty pcDNA3.1(+) was used to normalize DNA concentration. After 24h of transfection, cells were washed once with PBS, harvested in PBS, and spun down at 1,000g for 3 mn. Cell pellets were resuspended in 200 µl of phosphate lysis buffer (50 mM NaH2PO4, 150 mM NaCl, 1% Tween-20, 5% Glycerol, pH 8.0) supplemented with 2 µg/ml pepstatin, 2 µg/ml apoprotinin, 10 µg/ml leupeptin, 1 mM AEBSF [4-(2 Aminoethyl) benzenesulfonyl fluoride], and 1 mM Na3VO4. Lysis was performed on ice for ∼10 minutes with occasional vortexing, debris were centrifuged at 14,000 rpm for 10 mn before and protein concentration was calculated using Bradford reagent. For each condition, 20 to 30 µg of total protein was separated by SDS-PAGE on a pre-cast 4–15% Criterion™ Protein Gel (Bio-Rad Laboratories, Cat#5671084) followed by an overnight transfer and western blot analysis.

### In vitro charging assay of DUBs with the ISG15-VS activity-based probe

Charging of USP18, USP41, Ubp43, and all other variants, with the ISG15-VS activity-based probe (ABP) was done in a semi-pure way where DUBs were first expressed and immunoprecipitated from HEK-293T cells then used for in vitro assays. Typically, a 60 mm dish of confluent HEK-293T cells was transfected as described using 2.5 µg of DNA. After 24 to 48 hours of expression, cells were washed once with PBS, harvested in 1 ml of PBS then centrifuged at 1,000g for 3 mn, and cells pellets were either stored at -80°C or lysed immediately. Cell lysis was done using phosphate lysis buffer supplemented with 2 µg/ml pepstatin, 1 mM AEBSF [4-(2 Aminoethyl) benzenesulfonyl fluoride], 1 mM Na3VO4 and 10 mM DTT. Cell lysis was done for 10 mn on ice with occasional vortexing, then cell debris were removed by centrifugation at 14,000 rpm for 10 mn at 4°C. For each plate of cells expressing the desired construct, the DUB of interest was immunoprecipitated using 30 µl of anti-HA or anti-Myc beads for 1h at 4°C, then beads were washed once with lysis buffer (4°C), once with PBS (RT), resuspended in DUB activation buffer (pre warmed at 37°C) and split into 2 tubes. Beads were spun down, resuspended in 10 µl of DUB activation buffer (25 mM Tris pH 7.5, 150 mM NaCl and 10 mM DTT, pre warmed at 37°C), incubated for 10 mn at 37°C then mixed at a 1:1 ratio with DUB activation buffer containing 10 µM of ISG15-VS or no ABP as a negative control (buffer only). Reactions were incubated for 2 hours at 37°C, quenched by adding 10 µl of 4X Laemmli buffer buffer (62.5 mM Tris pH 6.7, 2% SDS, 10% glycerol, 100 mM DTT and 0.1% Bromophenol Blue) and separated by SDS-PAGE. Charging of DUBs with ISG15-VS was detected by western blotting the tag fused to the DUB.

### In vitro DUB assays using the fluorogenic substrate ISG15-AMC

The assay was conducted in a very similar way to the charging assay with ISG15-VS. Briefly, the indicated DUB constructs were expressed and purified from HEK-293T cells exactly as described above. Following IP of the DUBs, beads were washed once with lysis buffer (4°C), once with PBS (RT), and once with DUB buffer (50 mM Tris pH 7.5, 50 mM NaCl and 10 mM DTT, pre warmed at 37°C). Following the last wash, beads were resuspended in 40 µl of DUB buffer then split in 3 wells of a 384-well white flat plate (Corning, Cat# 3826BC), with 10 µl of immunoprecipitates per well. Following a 10 mn incubation at RT, each well was mixed with 10 µl of ISG15-AMC (R&D Systems, Cat# UL-553) that was diluted to 2 µM in DUB buffer. Release of AMC fluorescence was monitored using a BioTek Cytation 5 (Agilent Technologies) with an excitation wavelength of 380 nm and an emission wavelength of 460 nm. Reactions were carried out for either 45 or 60 mn, and readings were done every mn. Raw measurements were collected and presented as plots that were generated using GraphPad Prism.

### Generation of FAT10 conjugation in cells

To generate and analyze FAT10-conjugated proteins, we used an overexpression approach in HEK-293 cells based on a previously described protocol^44^. HEK-293 cells were first seeded in 10 cm dishes at approximately 2 million cells per dish. The day after, cells were transfected with 2 µg of 6HF-FAT10, either alone or in combination with 3 µg of HA-USP41 WT or 2CS, using PolyJet (SignaGen Laboratories, Cat# SL100688) and following the manufacturer’s instructions, except that the medium containing the transfection mix was replaced by fresh medium after 6 to 8 hours. After 48 hours of transfection, HEK-293 cells were washed with PBS a couple of times then harvested in PBS and centrifuged at 1,000g for 3 mn at 4°C. Cell pellets were resuspended in NETN buffer supplemented with protease inhibitors, then cell debris were removed by centrifugation at 14,000 rpm for 10 min at 4°C. Protein concentration was determined using Bradford reagent and inputs were prepared with the clarified lysate. The remaining lysate was mixed with anti-FLAG beads (20 µl per IP, Millipore Sigma) that were pre-washed with lysis buffer, and incubated on a rotary shaker for 2h at 4°C. Beads were washed with lysis buffer four times, resuspended in 25 µl of 2X Laemmli sample buffer, then inputs and IP samples were incubated at 95°C for 5 minutes before SDS-PAGE and western blot analysis.

### Structure generation using AlphaFold 3

We took advantage of AlphaFold 3 (https://alphafoldserver.com) to generate and predict the structures^40^ of USP18:hISG15 and USP41:FAT10. The sequences that were used can be accessed in the Supplementary Document 1 attached to this manuscript. In both cases, the top-ranked structures were selected then visualized using PyMOL. The PBD identifiers of the hISG15:PLpro^CoV2^ and mISG15:Ubp43 complexes solved by X-ray crystallography are 7RBS and 5CHV, respectively.

## AUTHOR CONTRIBUTIONS

TB and MJE conceptualized the project, conceived experiments, and analyzed data. TB carried out all experiments, except for the protein identification of USP18 Coomassie-stained bands by mass spectrometry, which was performed by the UNC Proteomics Core Facility. DLB and NGB assisted with generating AlphaFold 3 models and interpreting structural data. TB initially assembled figures and wrote the first draft of the manuscript. TB and MJE edited the manuscript and figures.

## ACKNOWLEDGEMENTS

Special thanks to the Emanuele laboratory for feedback and helpful discussions throughout this study. This work was supported by start-up funds from the University Cancer Research Fund (UCRF) to MJE, as well as grants from the National Institutes of Health (R01GM134231, R35GM153250), and the American Cancer Society (Research Scholar Grant; RSG-18-220-01-TBG). We thank Laura Herring, Natalie Barker, Thomas Webb and Angie Mordant, for their help with the proteomics work performed in this manuscript. This research is based in part upon work conducted using the UNC Proteomics Core Facility, which is supported in part by P30 CA016086 Cancer Center Core Support Grant to the UNC Lineberger Comprehensive Cancer Center.

---

**Supplementary Figure 1.**
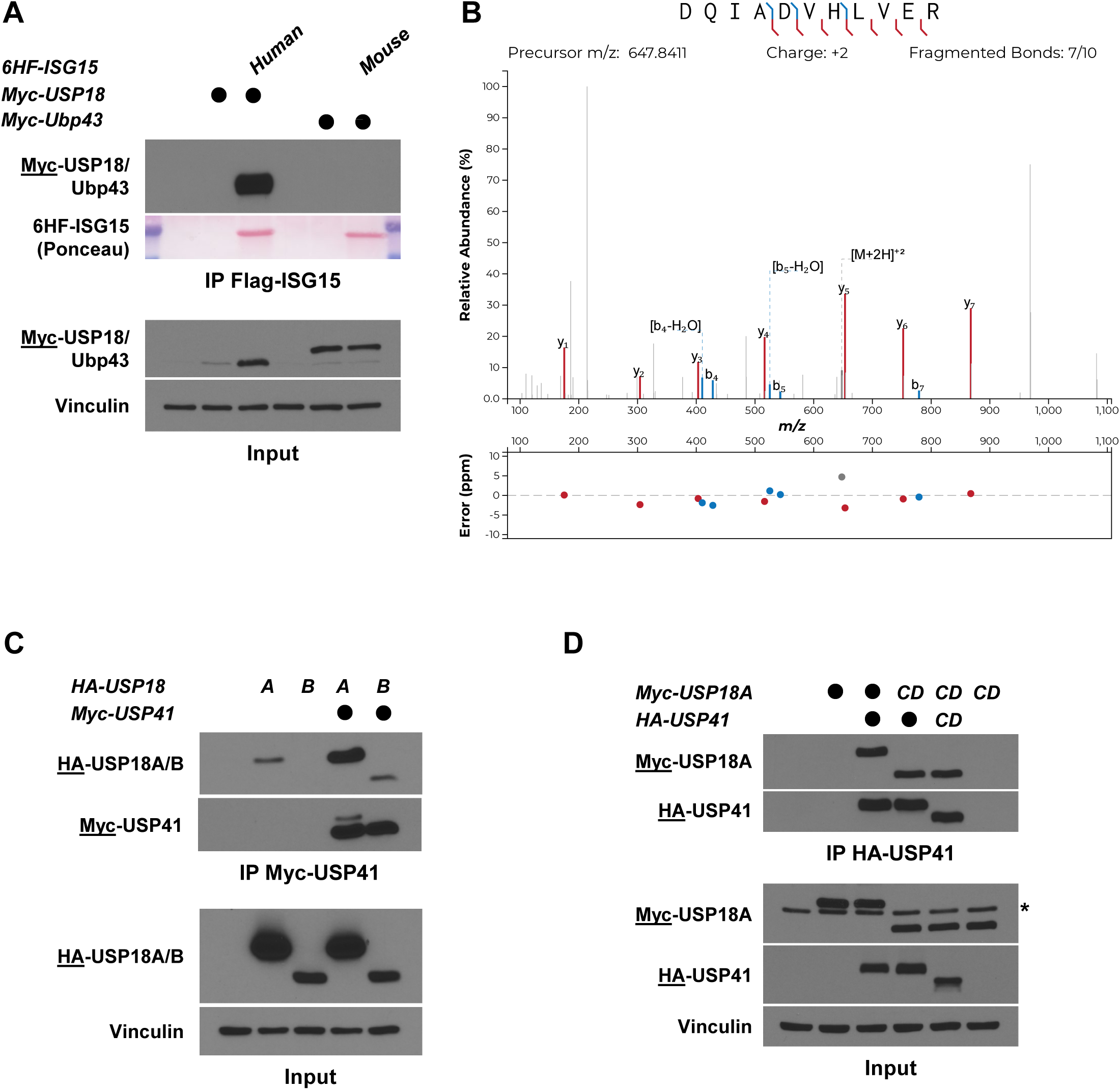
USP18 interacts with USP41. (a) Myc-USP18, or its mouse ortholog Myc-Ubp43, were ectopically expressed in HEK-293T cells alone or with human or mouse ISG15, respectively. After 24h, precleared lysates of transfected cells were subjected to immunoprecipitation on anti-FLAG beads. IP and input samples were separated by SDS-PAGE and analyzed by western blotting. Immunoblotted antigen is underlined to the left of blots. (b) MS/MS spectrum of the doubly charged ion (m/z 647.8411) corresponding to USP41 tryptic peptide DQIADVHLVER. (c) Myc-USP41 and HA-USP18 (both isoforms) were ectopically expressed in HEK-293T cells. After 24h, precleared lysates of transfected cells were used to immunoprecipitate USP41 on anti-Myc beads. Immunoblotted antigen is underlined to the left of blots. (d) Same as in (c) except that USP18A was used, as well as the catalytic domains (CD) of both USP41 and USP18A (USP41 CD and USP18A CD, respectively). Asterisk indicates a non-specific band reacting with the Myc antibody.

**Supplementary Figure 2.**
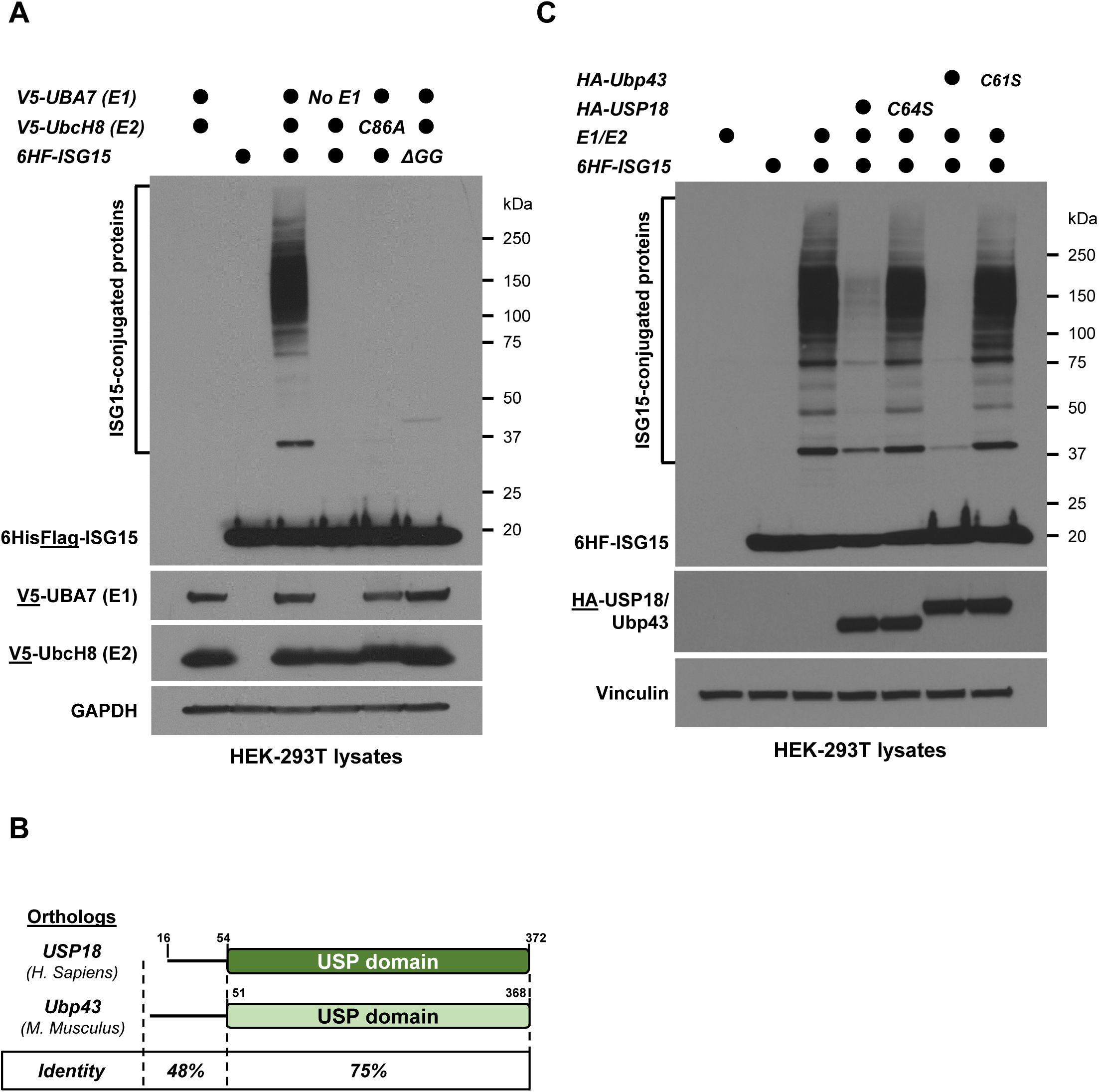
Overexpression system in HEK-293T to study protein ISGylation. (a) HEK-293T cells were transiently transfected with 6His-FLAG-ISG15 (6HF-ISG15) in combination with vectors encoding the ISG15 E1 activating enzyme (V5-UBA7) and the ISG15 E2 conjugating enzyme (V5-UbcH8). As negative controls, no E1, a catalytically inactive from of UbcH8 (C86A) or a non-conjugatable form of ISG15 (ΔGG) were used. Cell lysates were prepared 24 hours later and protein ISGylation was analyzed by western blotting using the indicated antibodies. (b) Cartoon showing the organization and sequence identity of USP18 and its mouse ortholog mUSP18/Ubp43. (c) Same as in Figure 2B, except that WT or catalytically inactive (C61S) versions of HA-Ubp43 were also included.

**Supplementary Figure 3.**
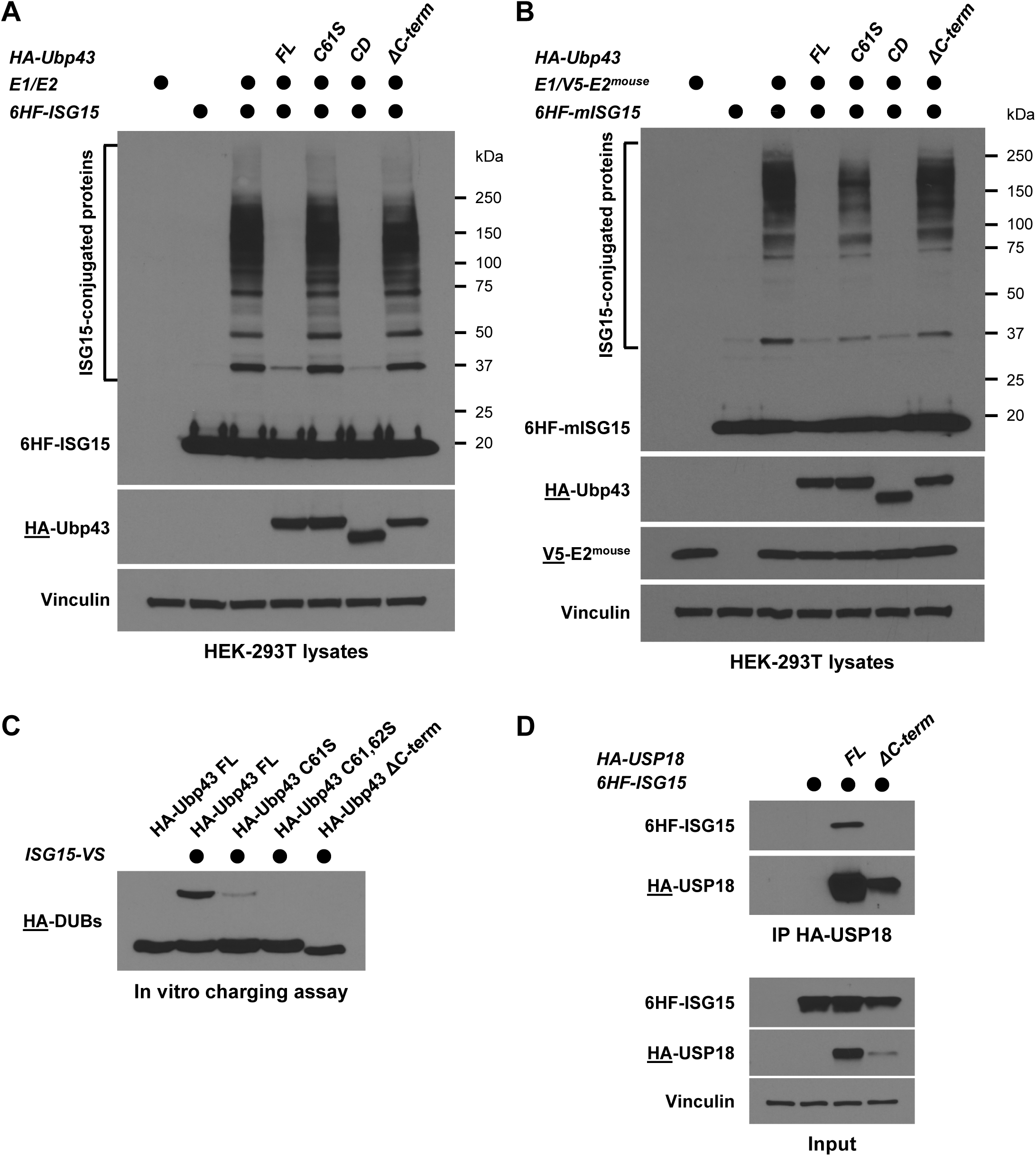
The C-terminus of Ubp43 is also necessary for its enzymatic activity. (a) Protein ISGylation in HEK-293T cells was reconstituted by transfection of the ISG15 machinery (E1/E2/ISG15) and where indicated, HA-tagged Ubp43 constructs were co-transfected. After 24h, lysates of transfected cells were prepared then analyzed by SDS-PAGE and western blot. Immunoblotted antigen is underlined to the left of blots. (b) Same as in (A), except that the mice version of both UbcH8 (E2^mouse^) and ISG15 (mISG15) were used. (c) The indicated HA-tagged Ubp43 constructs were ectopically expressed in HEK-293T cells and after 24h, precleared lysates of transfected cells were used to immunoprecipitate Ubp43 on anti-HA beads. Immunoprecipitates were mixed with reaction buffer containing ISG15-VS or not, and reaction products were analyzed by SDS-PAGE and western blot. (d) 6HF-ISG15 was ectopically expressed in HEK-293T cells, either alone or with HA-USP18 FL or HA-USP18 ΔC-term. After 24h, precleared lysates of transfected cells were used to immunoprecipitate USP18 FL or ΔC-term on anti-HA beads. Immunoblotted antigen is underlined to the left of blots.

**Supplementary Figure 4.**
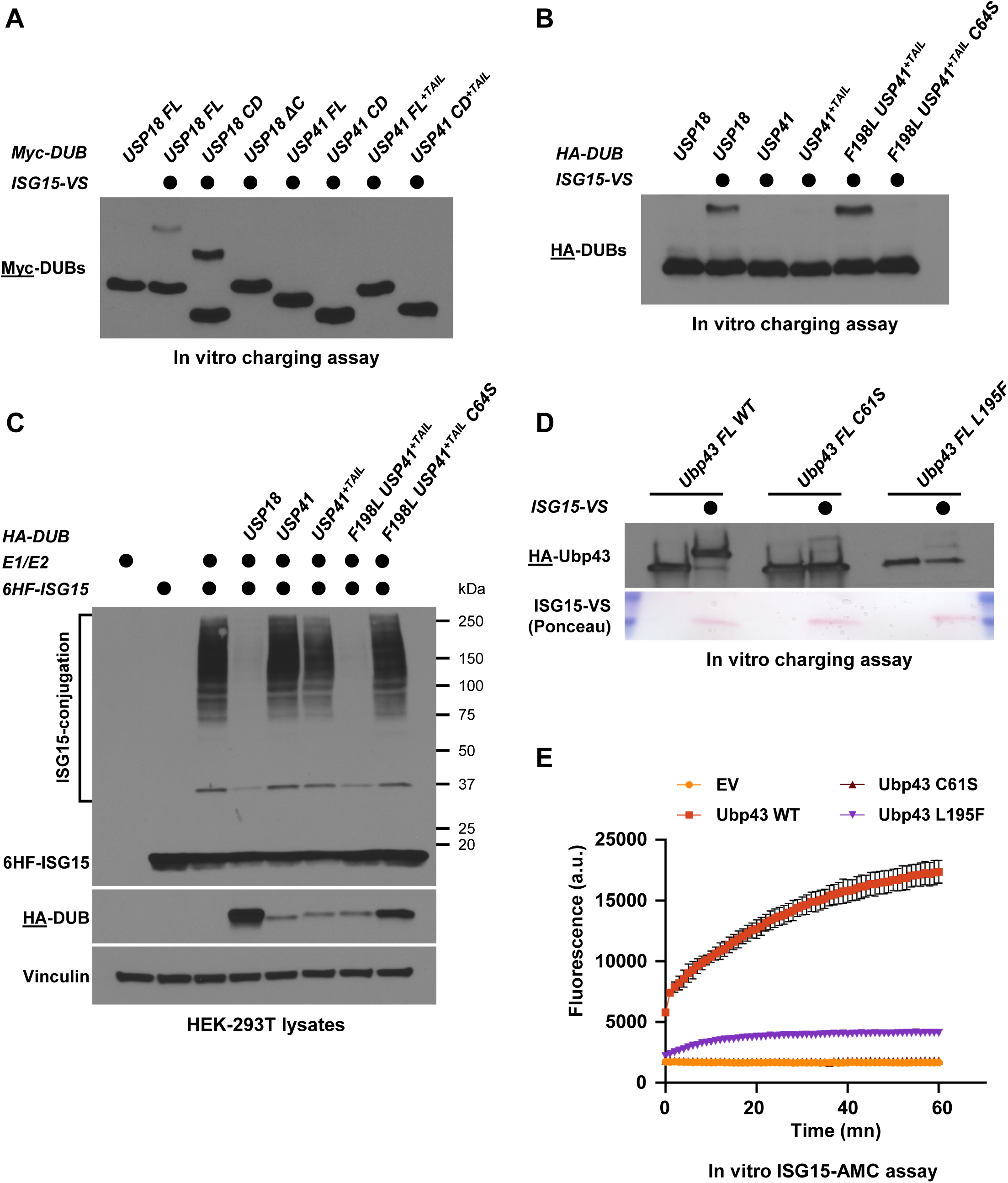
Leu198 is important for USP18 and mUSP18/Ubp43 catalytic activity. (a) The indicated Myc-tagged constructs were ectopically expressed in HEK-293T cells and after 24h, precleared lysates of transfected cells were used to immunoprecipitate these Myc-DUBs on anti-Myc beads. Immunoprecipitates were mixed with reaction buffer containing ISG15-VS, and reaction products were analyzed by SDS-PAGE and western blot. (b) Same as in (a), except that HA-tagged constructs were used. (c) Protein ISGylation in HEK-293T cells was reconstituted by transfection of the ISG15 machinery (E1/E2/ISG15) and where indicated, HA-tagged constructs were co-transfected. After 24h, lysates of transfected cells were prepared then analyzed by SDS-PAGE and western blot. Immunoblotted antigen is underlined to the left of blots. (d) The indicated HA-tagged versions of Ubp43 were isolated from HEK-293T cells on anti-HA beads 24h after transfection. HA-Ubp43 Immunoprecipitates were mixed with reaction buffer containing ISG15-VS or not, and reaction products were analyzed by SDS-PAGE and western blot. (e) The indicated HA-tagged constructs were ectopically expressed in HEK-293T cells and after 24h, precleared lysates of transfected cells were used to immunoprecipitate HA-tagged Ubp43 variants on anti-HA beads. Immunoprecipitates were mixed with reaction buffer containing the fluorogenic substrate ISG15-AMC, and fluorescence increase was monitored as previously described.

**Supplementary Figure 5.**
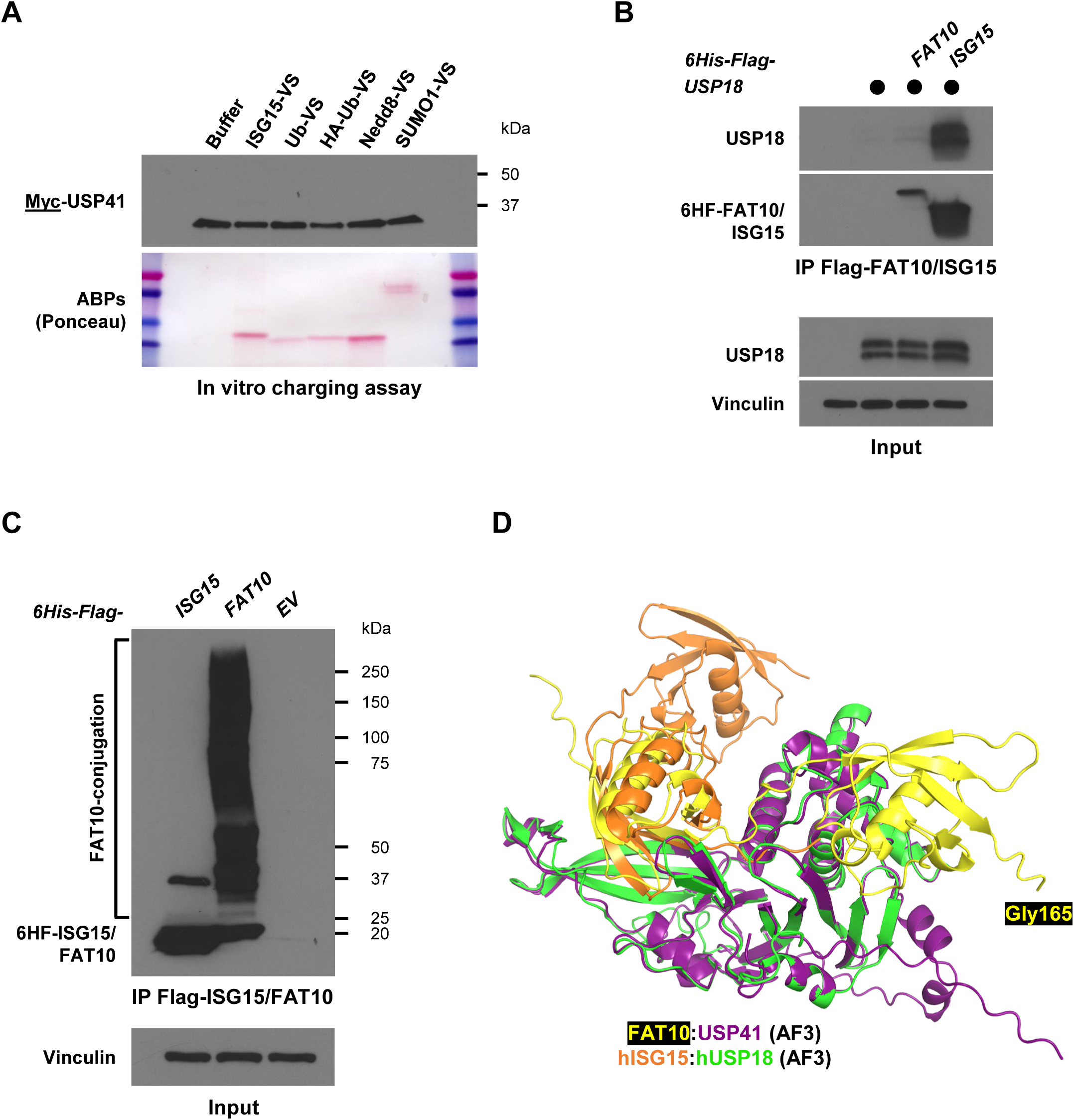
USP18 and USP41 interact with different Ubls. (a) Myc-USP41 was ectopically expressed in HEK-293T cells and after 24h, precleared lysates of transfected cells were used to immunoprecipitate USP41 on anti-Myc beads. Immunoprecipitates were mixed with reaction buffer containing the indicated activity-based probes (ABPs), and reaction products were analyzed by SDS-PAGE and western blot. (b) Untagged USP18 was ectopically expressed in HEK-293 cells, either alone or with 6HF-FAT10 or 6HF-ISG15. After 24h, precleared lysates of transfected cells were used to immunoprecipitate FAT10 and ISG15 on anti-FLAG beads. Immunoprecipitates and inputs were analyzed by western blotting. (c) 6HF-ISG15 or 6HF-FAT10 were ectopically expressed in HEK-293 cells. After 48h, precleared lysates of transfected cells were used to immunoprecipitate FAT10-conjugated proteins on anti-FLAG beads. IP and Input samples were analyzed by SDS-PAGE and western blot. (d) Overlay of FAT10:USP41 with hUSP18:hISG15 by aligning the structures on the DUBs.

**Supplementary Figure 6.**
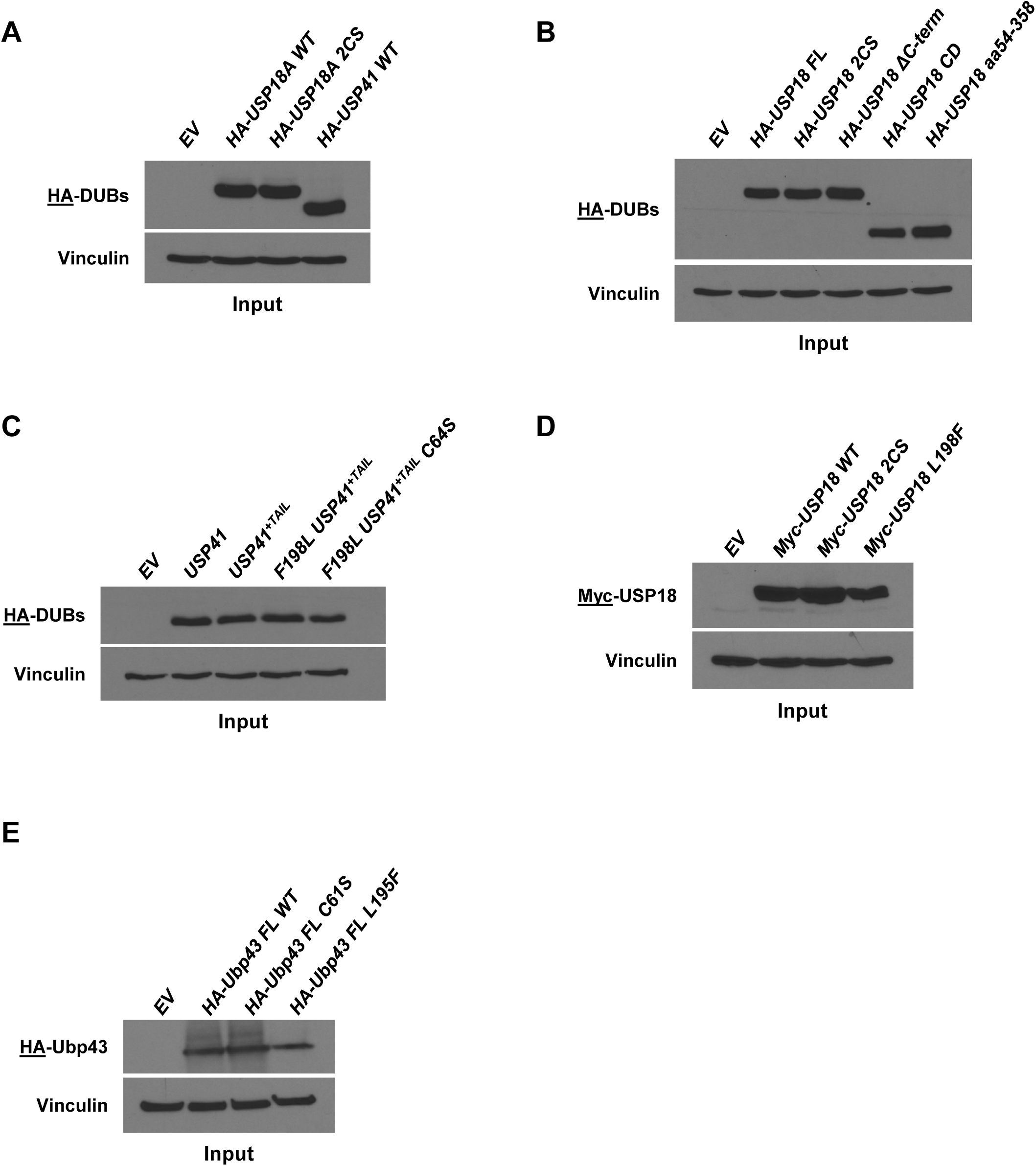
Control of expression for ISG15-AMC experiments. (a) HEK-293T lysates from cells transfected with the indicated constructs were separated by SDS-PAGE and analyzed by western blotting using the indicated antibodies. This serves as a control for the experiment presented in Figure 2D. (b) Cell lysates of HEK-293T transfected with the indicated HA-tagged USP18 constructs were separated by SDS-PAGE and analyzed by western blotting. This serves as a control for the experiment presented in Figure 3C. (c) Cell lysates of HEK-293T transfected with the indicated HA-tagged USP18 constructs were separated by SDS-PAGE and analyzed by western blotting. This serves as a control for the experiment presented in Figure 4D. (d) Cell lysates of HEK-293T transfected with the indicated Myc-tagged USP18 constructs were separated by SDS-PAGE and analyzed by western blotting. This serves as a control for the experiment presented in Figure 4F. (e) Cell lysates of HEK-293T transfected with the indicated HA-tagged Ubp43 constructs were separated by SDS-PAGE and analyzed by western blotting. This serves as a control for the experiment presented in Supplementary Figure 4E.

